# Integrating deep learning and unbiased automated high-content screening to identify complex disease signatures in human fibroblasts

**DOI:** 10.1101/2020.11.13.380576

**Authors:** Lauren Schiff, Bianca Migliori, Ye Chen, Deidre Carter, Caitlyn Bonilla, Jenna Hall, Minjie Fan, Edmund Tam, Sara Ahadi, Brodie Fischbacher, Anton Geraschenko, Christopher J. Hunter, Subhashini Venugopalan, Sean DesMarteau, Arunachalam Narayanaswamy, Selwyn Jacob, Zan Armstrong, Peter Ferrarotto, Brian Williams, Geoff Buckley-Herd, Jon Hazard, Jordan Goldberg, Marc Coram, Reid Otto, Edward A. Baltz, Laura Andres-Martin, Orion Pritchard, Alyssa Duren-Lubanski, Ameya Daigavane, Kathryn Reggio, NYSCF Global Stem Cell Array^®^ Team, Phillip C. Nelson, Michael Frumkin, Susan L. Solomon, Lauren Bauer, Raeka S. Aiyar, Elizabeth Schwarzbach, Scott A. Noggle, Frederick J. Monsma, Daniel Paull, Marc Berndl, Samuel J. Yang, Bjarki Johannesson

## Abstract

Drug discovery for diseases such as Parkinson’s disease are impeded by the lack of screenable cellular phenotypes. We present an unbiased phenotypic profiling platform that combines automated cell culture, high-content imaging, Cell Painting, and deep learning. We applied this platform to primary fibroblasts from 91 Parkinson’s disease patients and matched healthy controls, creating the largest publicly available Cell Painting image dataset to date at 48 terabytes. We use fixed weights from a convolutional deep neural network trained on ImageNet to generate deep embeddings from each image and train machine learning models to detect morphological disease phenotypes. Our platform’s robustness and sensitivity allow the detection of individual-specific variation with high fidelity across batches and plate layouts. Lastly, our models confidently separate *LRRK2* and sporadic Parkinson’s disease lines from healthy controls (receiver operating characteristic area under curve 0.79 (0.08 standard deviation)), supporting the capacity of this platform for complex disease modeling and drug screening applications.

---

A major challenge in discovering effective therapies for complex diseases is defining robust disease phenotypes amenable to high-throughput drug screens^1^. The increasing availability of patient cells through biobanking and induced pluripotent stem cell (iPSC) models presents an excellent opportunity for cell-based drug discovery, but in the absence of reliable drug targets, new methods to discover unbiased, quantitative cellular phenotypes are still needed. Recent advancements in artificial intelligence (AI) and deep learning–based analysis offer the potential to accelerate therapeutic discovery by distinguishing drug-induced cellular phenotypes^2^, elucidating mechanisms of action^3^, and gaining insights into drug repurposing^4,5^. Applying similar unbiased approaches to large, high-quality datasets using methods such as high-content imaging is a powerful strategy to capture novel patient- or disease-specific phenotypic patterns. Several studies have applied AI to large datasets to uncover population-based disease phenotypes and biomarkers, but the power of these studies thus far has been limited by small cohort sizes^6,7^ and non-physiological cellular perturbations^8,9^.

Parkinson’s disease (PD) is the second most prevalent progressive neurodegenerative disease, affecting 2–3% of individuals over the age of 65^10^. While variants in many genes including *LRRK2^11^*, *GBA^12^*, and *SNCA^13^* have been associated with disease risk, over 90% of cases are sporadic, caused by unknown genetic and environmental factors^14^. Substantial progress has been made in elucidating the pathological mechanisms underlying PD, but the failure of recent clinical trials targeting established pathological pathways suggests that current drug discovery strategies remain inadequate^15^. This challenge is exacerbated by the lack of animal models that sufficiently recapitulate PD pathology^16^ and of cellular phenotypes amenable to drug screening. Though it is known that PD pathogenesis involves a complex concert of events and molecular players, current strategies for drug discovery largely focus on hypothesis-driven approaches where single, predefined cellular readouts or genetic variants are targeted^17,18^.

In this study, we combined scalable automation and deep learning to develop a high-throughput and high-content screening platform for unbiased population-scale morphological profiling of cellular phenotypes. We applied this platform to primary PD fibroblasts, a readily accessible cell type that reflects donor genetics and environmental exposure history^7,19^. Our highly standardized automation procedures allowed for model generalization across batches, demonstrated by the power of our platform to recognize individual cell lines within a pool of 96 lines, even across batches and plate layouts. Furthermore, cells acquired from multiple biopsies from the same individual, but collected years apart, resulted in more similar morphological profiles than cells derived from different individuals. Importantly, our unbiased profiling approach also identified generalizable PD disease signatures, which allowed us to distinguish both sporadic PD and *LRRK2* PD cells from those of healthy controls. Taken together, our deep learning–based platform provides a powerful approach for de novo, unbiased identification of quantitative cellular disease phenotypes that can be leveraged for drug screening.

## Results

### Automated high-content phenotyping platform achieves high data consistency

We developed an automated platform to morphologically profile large collections of cells leveraging the capabilities of the New York Stem Cell Foundation (NYSCF) Global Stem Cell Array^®^, a modular robotic platform for large-scale cell culture automation^20^, and applied it to search for Parkinson’s disease-specific cellular signatures in primary human fibroblasts (**Fig. 1a**). Starting from a collection of more than 1000 fibroblast lines in the NYSCF repository that were collected and derived using highly standardized methods^20^, we selected a subset of PD lines from sporadic patients and patients carrying *LRRK2* (G2019S) or *GBA* (N370S) mutations, as well as age-, sex-, and ethnicity-matched (based on self-identification) healthy controls. All lines underwent thorough genetic quality control and exclusion criteria–based profiling, which yielded lines from 45 healthy controls, 32 sporadic PD, 8 *GBA* PD and 6 *LRRK2* PD donors; 5 participants also donated a second skin biopsy 3 to 6 years later, which were analyzed as independent lines, for a total of 96 cell lines (Methods, **Supplementary Data 1**).

**Fig. 1.**
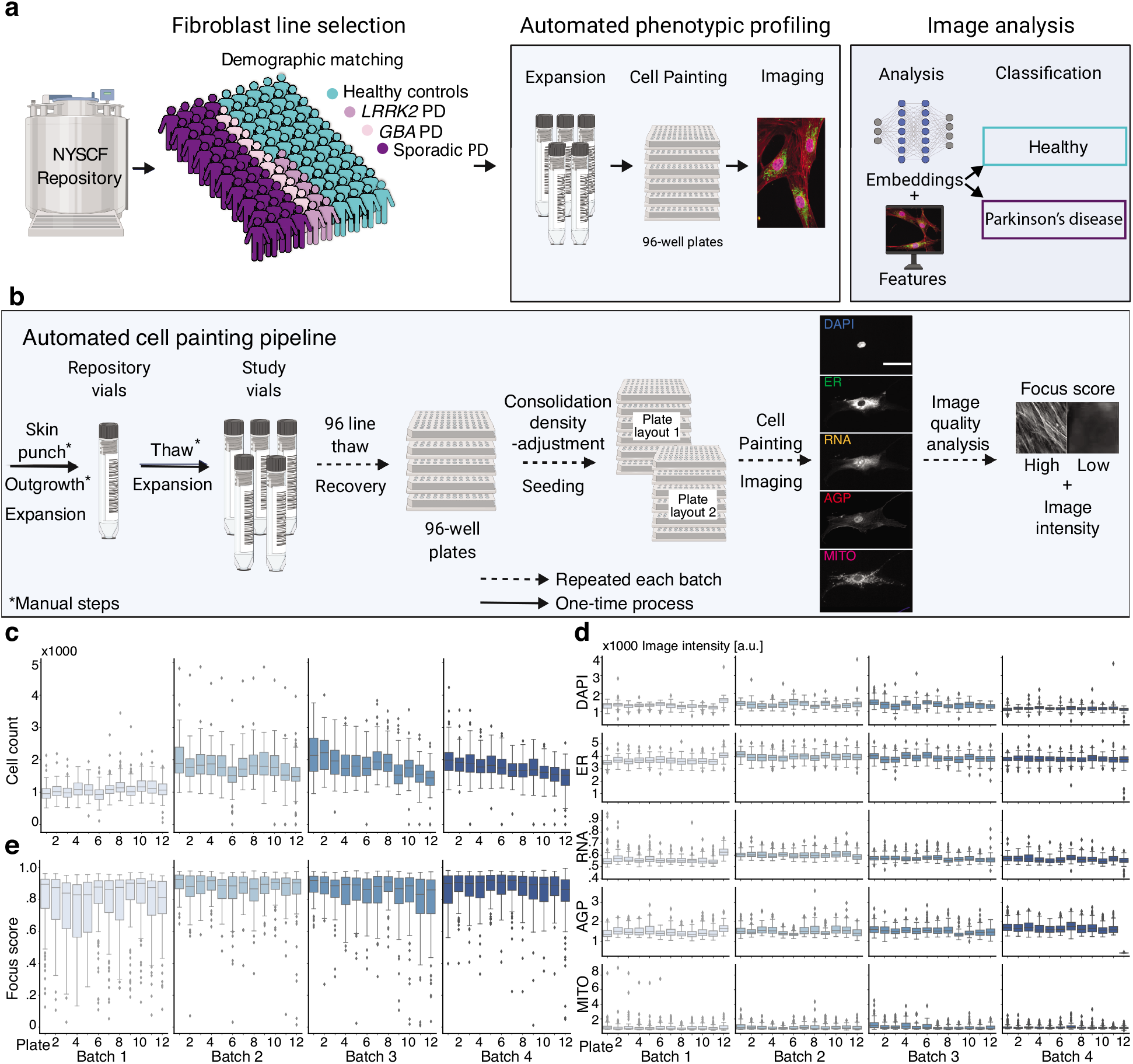
Automated high-content profiling platform demonstrates reproducibility across batches. **a**, Workflow overview. **b**, Overview of automated experimental pipeline. Scale bar: 35 μm. Running this pipeline yielded low variation across batches in: (**c**) well-level cell count; (**d**) well-level foreground staining intensity distribution per channel and plate; and (**e**) well-level image focus across the endoplasmic reticulum (ER) channel per plate, for n=96 wells per plate. Box plot components are: horizontal line, median; box, interquartile range; whiskers, 1.5× interquartile range; black squares, outliers.

We then applied our automated procedures for cell thawing, expansion and seeding, which were designed to minimize experimental variation and maximize reproducibility across plates and batches (**Fig. 1b**). This method resulted in consistent growth rates across all 4 experimental groups during expansion (**Supplementary Fig. 1**), although some variation was seen in assay plate cell counts (**Fig. 1c**). Importantly, overall cell counts for healthy and PD cell lines remained highly similar (**Supplementary Fig. 1**).

Two days after seeding into assay plates, we applied automated procedures to stain the cells with Cell Painting dyes^21^ for multiplexed detection of cell compartments and morphological features (nucleus (DAPI), nucleoli and cytoplasmic RNA (RNA), endoplasmic reticulum (ER), actin, golgi and plasma membrane (AGP), and mitochondria (MITO)). Plates were then imaged in 5 fluorescent channels with 76 tiles per well, resulting in uniform image intensity and focus quality across batches (**Fig. 1d, e**) and ~1 terabyte of data per plate. Additionally, to ensure consistent data quality across wells, plates and batches, we built an automated tool for near realtime quantitative evaluation of image focus and staining intensity within each channel (**Supplementary Fig. 2**). The tool is based on random sub-sampling of tile images within each well of a plate to facilitate immediate analysis and has been made publicly available (see Code Availability statement). Finally, the provenance of all but two cell lines were confirmed (see Methods). In summary, using scalable automation, we built an end-to-end platform that consistently and robustly thaws, expands, plates, stains, and images primary human fibroblasts for phenotypic screening.

### Experimental strategy for achieving unbiased deep learning-based image analysis

To analyze our high-content imaging data, we built a custom unbiased deep learning pipeline. In our pipeline, both cropped cell images and tile images (full-resolution microscope images) were fed through an Inception architecture deep convolutional neural network^22^ that had been pretrained on ImageNet, an object recognition dataset^23^, to generate deep embeddings that could be viewed as lower-dimensional morphological profiles of the original images (**Fig. 2a**). In this dataset, each tile or cell was represented as a 64-dimensional vector for each of the 5 fluorescent channels, which were combined into a 320-dimensional deep embedding vector.

**Fig. 2.**
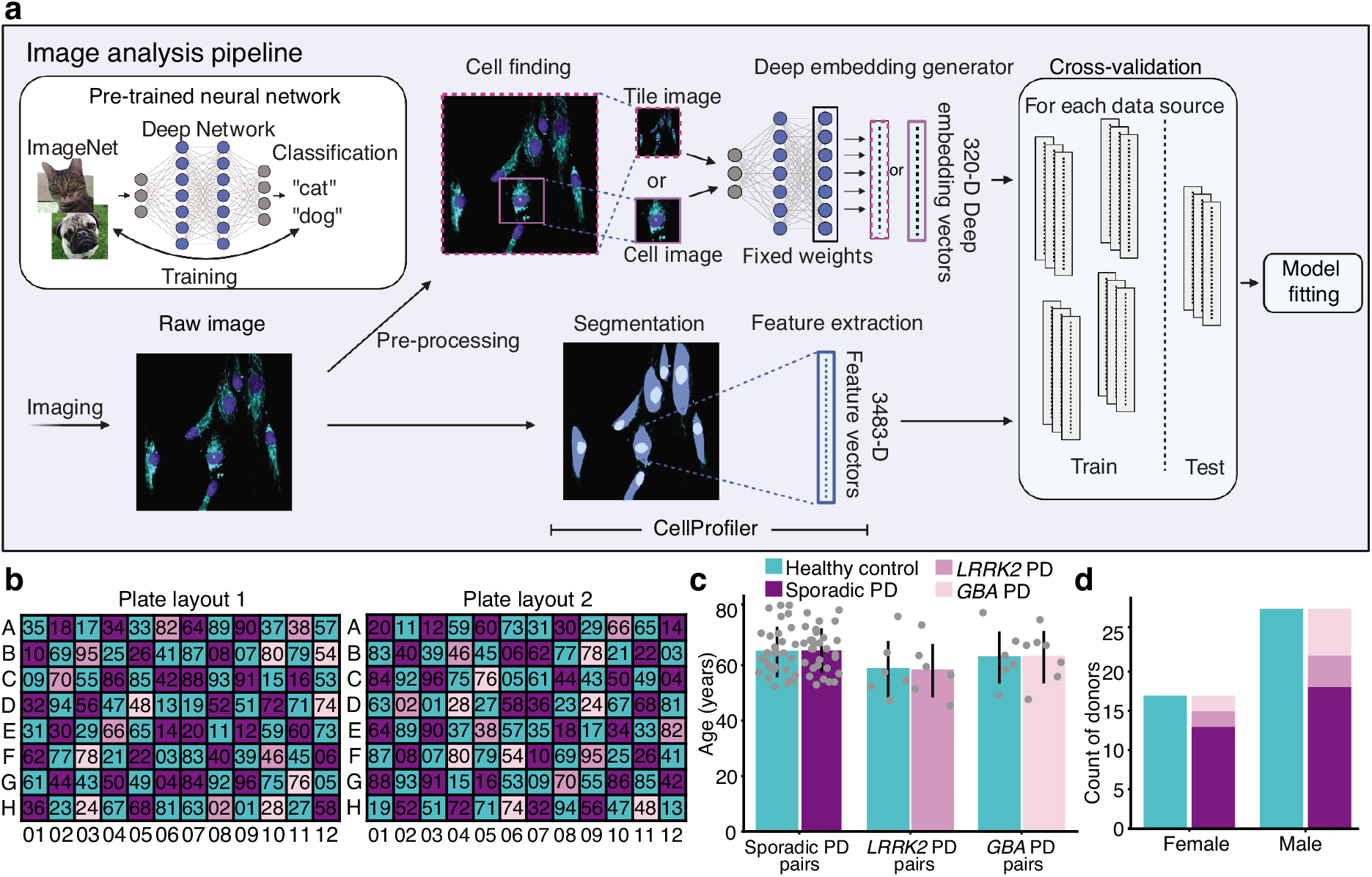
Image analysis pipeline and rigorous experimental design enable unbiased deep learning-based high-content screening. **a**, A deep embedding generator (a neural network pre-trained on an independent object recognition task) maps each tile or cell image independently to a deep embedding vector, which along with CellProfiler features and basic image statistics were used as data sources for model fitting and evaluation for supervised learning prediction tasks including healthy vs. PD classification. **b**, Two 96-well plate layouts used in each experimental batch control for location biases. In each layout, each well contained cells from one cell line denoted by the two-digit label. The second layout consisted of diagonally translating each of the four quadrants of the first. n=45 healthy controls and n=45 PD patients were matched in pairs based on demographics, including by (**c**) age and (**d**) sex. Data are presented as mean values +/− standard deviation.

For a more comprehensive analysis, we also used as a baseline basic image statistics (e.g. image intensity) and conventional cell image features extracted by a CellProfiler^24^ pipeline that computes 3483 features from each segmented cell. CellProfiler features, albeit potentially less accurate than deep image embeddings in some modeling tasks^2^, provide a comprehensive set of hand-engineered measurements that have a direct link to a phenotypic characteristic, facilitating biological interpretation of the phenotypes identified.

For modeling, we included several standard supervised machine learning models including random forest, multilayer perceptron and logistic regression classifier models, as well as ridge regression models. All of these models output a prediction based on model weights fitted to training data, but can have varying performance based on the structure of signal and noise in a given dataset. We trained these models on the well-average deep embedding and feature vectors. Specifically, we took the average along each deep embedding or feature dimension to obtain a single data point representative of all cellular phenotypes within a well. To appropriately assess model generalization on either data from new experiments or data from new individuals, we adopted k-fold cross-validation stratified by batch or individuals for cell line and disease prediction, respectively.

Since deep learning-based analysis is highly sensitive, including to experimental confounds, we ensured each 96-well plate contained all 96 cell lines (one line per well) and incorporated two distinct plate layout designs to control for potential location biases (**Fig. 2b**). The plate layouts alternate control and PD lines every other well and also position control and PD lines paired by both age (**Fig. 2c**) and sex (**Fig. 2d**) in adjacent wells, when possible. Importantly, we quantitatively confirmed the robustness of our experimental design by performing a lasso variable selection for healthy vs. PD on participant, cell line, and plate covariates, which did not reveal any significant biases (**Supplementary Fig. 1**). We conducted four identical batches of the experiment, each with six replicates of each plate layout, yielding 48 plates of data, or approximately 48 wells for each of the 96 cell lines. In summary, we employed a robust experimental design that successfully minimized the effect of potential covariates. We also established a comprehensive image analysis pipeline where multiple machine learning models were applied to each classification task, using both computed deep embeddings and extracted cell features as data sources.

### Identification of individual cell lines based on morphological profiles using deep learning models

The strength and challenge of population-based profiling is the innate ability to capture individual variation. Similarly, the variation of high-content imaging data generated in separate batches is also a known confound in large-scale studies. Evaluating a large number of compounds, or, in this case, a large number of replicates to achieve a sufficiently strong disease model, necessitates aggregating data across multiple experimental batches. We sought to assess the line-to-line and batch-to-batch variation in our dataset by evaluating if a model trained to identify an individual cell line could successfully identify that same cell line in an unseen batch among *n* = 96 cell lines (**Fig. 3a**), because the inability to do so might make establishing a consistent disease model across experimental batches challenging. To this end, we adopted a 4-fold cross-validation scheme, where a model was fit to three out of four batches and its performance assessed with accuracy as a metric (number of correct predictions divided by total examples) was evaluated on the fourth, held-out batch. Importantly, we also held out the plate layout to ensure that the model was unable to rely on any possible location biases (**Supplementary Table 1**).

**Fig. 3.**
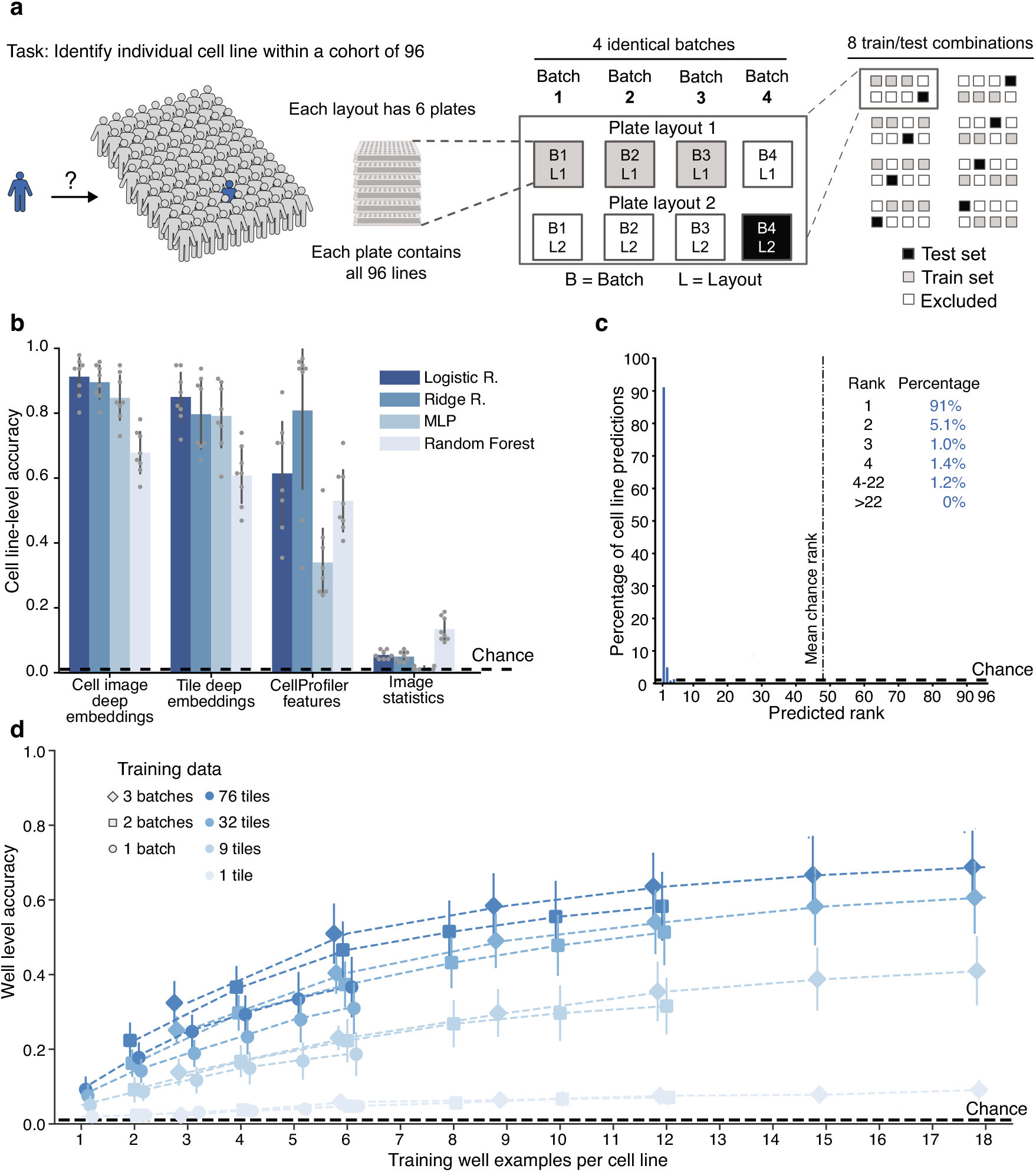
Robust identification of individual cell lines across batches and plate layouts. **a**, 96-way cell line classification task uses a cross-validation strategy with held-out batch and plate-layout. **b**, Test set cell line–level classification accuracy is much higher than chance for both deep image embeddings and CellProfiler features using a variety of models (logistic regression, ridge regression, multilayer perceptron (MLP), and random forest). Error bars denote standard deviation across 8 batch/plate layouts. **c**, Histogram of cell line–level predicted rank of true cell line for the logistic regression model trained on cell image deep embeddings from **b** shows that the correct cell line is ranked first in 91% of cases. **d**, A multilayer perceptron model trained on smaller cross sections of the entire dataset, down to a single well (average of cell image deep embeddings across 76 tiles) per cell line, can identify a cell line in a held-out batch and plate layout with higher than chance well-level accuracy; accuracy rises with increasing training data. Data are presented as mean values +/− standard deviation. Dashed black lines denote chance performance.

Our analysis revealed that models trained on CellProfiler features and deep image embeddings performed better than chance and the baseline image statistics (**Fig. 3b**). The logistic regression model trained on well-mean cell image deep embeddings (a single 320-D vector representing each well) achieved a cell line–level (average predictions across all six held-out test wells) accuracy of 91% (6% SD), compared to a 1.0% (1 out of 96) expected accuracy by chance alone. In cases when this model’s prediction was incorrect, the predicted rank of the correct cell line was still at most within the top 22 out of 96 lines (**Fig. 3c**). A review of the model’s errors presented as a confusion matrix did not reveal any particular pattern in the errors (**Supplementary Fig. 3**). In summary, our results show that our model can successfully detect variation between individual cell lines by correctly identifying cell lines across different experimental batches and plate layouts.

We then set out to determine how the quantity of available training data impacts the detection of this cell line–specific signal (**Fig. 3d**). We varied the training data by reducing the number of tile images per well (from 76 to 1) and well examples (from 18 to 1 (6 plates per batch and 3 batches to 1 plate from 1 batch)) per cell line with a multilayer perceptron model (which can be trained on a single data point per class). We trained on well-averaged cell image deep embeddings and evaluated on a held-out batch using well-level accuracy by taking only the prediction from each well, without averaging multiple such predictions. Although reducing the number of training wells per cell line or tiles per well reduced accuracy, remarkably, a model trained on just a single well data point (the average of cell image deep embeddings from 76 tiles in that well) per cell line from a single batch still achieved 9% (3% SD) accuracy, compared to 1.0% chance. Collectively, these results indicate the presence of robust line-specific signatures, which our deep learning platform is notably able to distinguish with minimal training data.

### Cell morphology is similar across multiple lines from the same donor

Next, we assessed whether the identified signal in a given cell line was in fact a characteristic of the donor rather than an artifact of the cell line handling process or biopsy procedures (e.g., location of skin biopsy). For this purpose, we leveraged the second biopsy samples provided by 5 of the 91 donors 3 to 6 years after their first donation. We retrained the logistic regression on cell image deep embeddings on a modified task consisting of only one cell line from each of the 91 donors with batch and plate layout held out as before (**Fig. 4a**, **Supplementary Table 2**). After training, we tested the model by evaluating the ranking of the 5 held-out second skin biopsies among all 91 possible predictions, in the held-out batch and plate-layout. This train and test procedure was repeated, interchanging whether the held-out set of lines corresponded to the first or second skin biopsy.

**Fig. 4.**
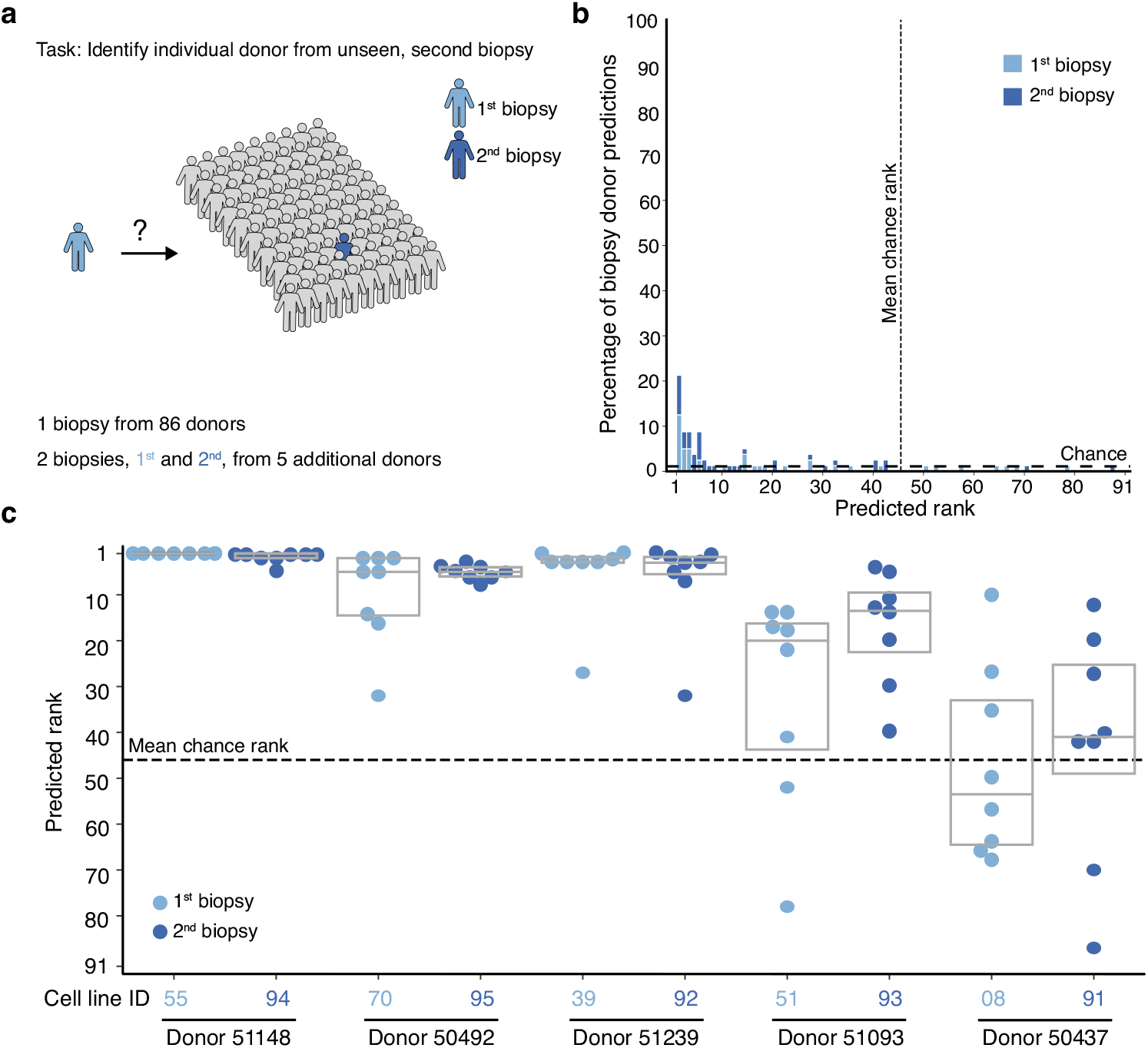
Donor-specific signatures revealed in analysis of repeated biopsies from individuals. **a**, The 91-way biopsy donor classification task uses a cross-validation strategy with held-out cell lines, and also held-out batch and plate layout as in **Fig. 3a**. **b**, Histogram and (**c**) box plots of test set cell line–level predicted rank among 91 biopsy donors of the 8 held-out batch/plate layouts for 10 biopsies (first and second from 5 individuals) assessed, showing the correct donor is identified in most cases for 4 of 5 donors. Dashed lines denote chance performance. Box plot components are: horizontal line, median; box, interquartile range.

The models achieved 21% (13% SD) accuracy in correctly identifying which of the 91 possible donors the held-out cell line came from, compared to 1.1% (1 out of 91) by chance (**Fig. 4b**). In cases where the model’s top prediction was incorrect, the predicted rank of the correct donor was much higher than chance for four of the five donors (**Fig. 4c**), even though the first and second skin biopsies were acquired years apart. In one case (donor 51239), the second biopsy was acquired from the right arm instead of the left arm, but the predicted rank was still higher than chance. The one individual (donor 50437) whose second biopsy was not consistently ranked higher than chance was the only individual who had one of the two biopsies acquired from the leg instead of both biopsies taken from the arm. Taken together, our model was able to identify donor-specific variations in morphological signatures that were unrelated to cell handling and derivation procedures, even across experimental batches.

### Deep learning-based morphological profiling can separate PD fibroblasts (sporadic and *LRRK2*) from healthy controls

Next, we evaluated the ability of our platform to achieve its primary goal of distinguishing between cell lines from PD patients and healthy controls. We divided sporadic PD, *LRRK2* PD participants, and paired demographically matched healthy controls (*n* = 74 participants) into 5 groups (**Supplementary Data 2**) for 5-fold cross-validation, where a model is trained to predict healthy or PD on 4 of the 5 sets of the cell line pairs and tested on the held-out 5th set of cell lines (**Fig. 5a**). To evaluate performance, we used the area under the receiver operating characteristic curve (ROC AUC) metric, which evaluates the probability of ranking a random healthy cell line as “more healthy” than a random PD cell line, where 0.5 ROC AUC is chance and 1.0 is a perfect classifier. Following training, we evaluated the ROC AUC on the test set in three ways: first with both sporadic and *LRRK2* PD (*n* = 37 participants) vs. all controls (*n* = 37 participants), then with the sporadic PD (*n* = 31 participants) vs. all controls (*n* = 37 participants), and then with *LRRK2* PD (*n* = 6 participants) vs. all controls (*n* = 37 participants). Finally, we ensured these results are robust to the cell line with a potentially unconfirmed disease state (Methods).

**Fig. 5.**
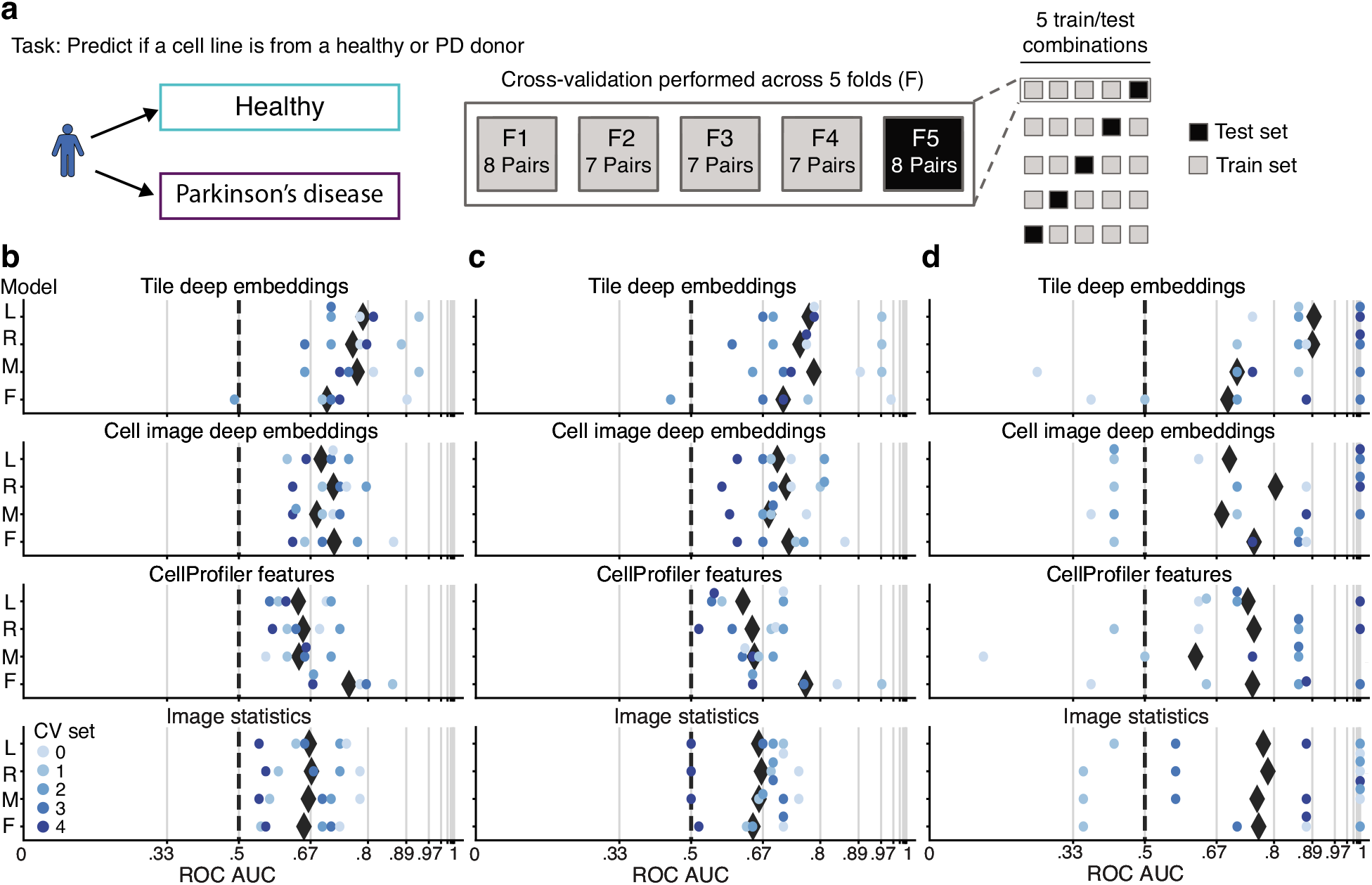
PD-specific signatures identified in sporadic and *LRRK2* PD primary fibroblasts. **a**, PD vs. healthy classification task uses a k-fold cross-validation strategy with held-out PD-control cell line pairs. Cell line–level ROC AUC, the probability of correctly ranking a random healthy control and PD cell line evaluated on held out–test cell lines for (**b**) *LRRK2/sporadic* PD and controls (**c**) sporadic PD and controls and (**d**) *LRRK2* PD and controls, for a variety of data sources and models (logistic regression (L), ridge regression (R), multilayer perceptron (M), and random forest (F)), range from 0.79–0.89 ROC AUC for the top tile deep embedding model and 0.75–0.77 ROC AUC for the top CellProfiler feature model. Black diamonds denote the mean across all cross-validation (CV) sets. Grid line spacing denotes a doubling of the odds of correctly ranking a random control and PD cell line and dashed lines denote chance performance.

We omitted *GBA* PD samples from this evaluation because our analysis of these lines indicated a possible underlying stratification imbalance^25^ in our partitioning across cross-validation datasets, demonstrated by a majority of the splits achieving ROC AUCs well below 0.5 (**Supplementary Fig. 4**). As a donor metadata review did not reveal a clear cause for this imbalance, we expect it might only be resolved with a larger, more representative set of *GBA* PD samples.

As in the above analyses, we used both cell and tile deep embeddings, CellProfiler features, and image statistics as data sources for model fitting in PD vs. healthy classification. The model with the highest mean ROC AUC, a logistic regression trained on tile deep embeddings, achieved a 0.79 (0.08 SD) ROC AUC for PD vs. healthy, while a random forest trained on CellProfiler features achieved a 0.76 (0.07 SD) ROC AUC (**Fig. 5b**). To investigate if the signal was predominantly driven by one of the PD subgroups, we probed the average ROC AUCs for each one. The model trained on tile deep embeddings achieved a 0.77 (0.10 SD) ROC AUC for separating sporadic PD from controls and 0.89 (0.10 SD) ROC AUC for separating *LRRK2* PD from controls (**Fig. 5c, d**), indicating that both patient groups contain strong diseasespecific signatures.

Finally, to investigate the source of the predictive signal, we studied the performance of the logistic regression trained on tile deep embeddings, but where the data either omitted one of the five Cell Painting stains or included only a single stain, in performing sporadic and *LRRK2* PD vs. healthy classification (**Supplementary Fig. 5**). Interestingly, the performance was only minimally affected by the removal of any one channel, indicating that the signal was robust. These results demonstrate that our platform can successfully distinguish PD fibroblasts (either *LRRK2* or sporadic) from control fibroblasts.

### Fixed feature extraction and analysis reveal biological complexity of PD-related signatures

Lastly, we further explored the CellProfiler features to investigate which biological factors might be driving the separation between disease and control, focusing on random forest, ridge regression, and logistic regression model architectures, as these provide a ranking of the most meaningful features. We first estimated the number of top-ranking features among the total set of 3483 features that were sufficient to retain the performance of the random forest classifier on the entire feature set and found the first 1200 to be sufficient (**Supplementary Fig. 6**).

Among the top 1200 features of each of the 3 model architectures (each with 5 crossvalidation folds), 100 features were present in all 15 folds (**Fig. 6a**). From among these, we removed correlated features using a Pearson correlation threshold of 0.75, leaving 55 uncorrelated features (**Supplementary Table 3**). To see if these best performing features held any mechanistic clues, we grouped them based on their type of measurement (e.g., shape, texture and intensity) and their origin by cellular compartment (cell, nucleus or cytoplasm) or image channel (DAPI, ER, RNA, AGP, and MITO)^26^. Such groupings resulted in features implicated in “area and shape,” “radial distribution” of signal within the RNA and AGP channels, and the “granularity” of signal in the mitochondria channel (**Fig. 6b**).

**Fig. 6.**
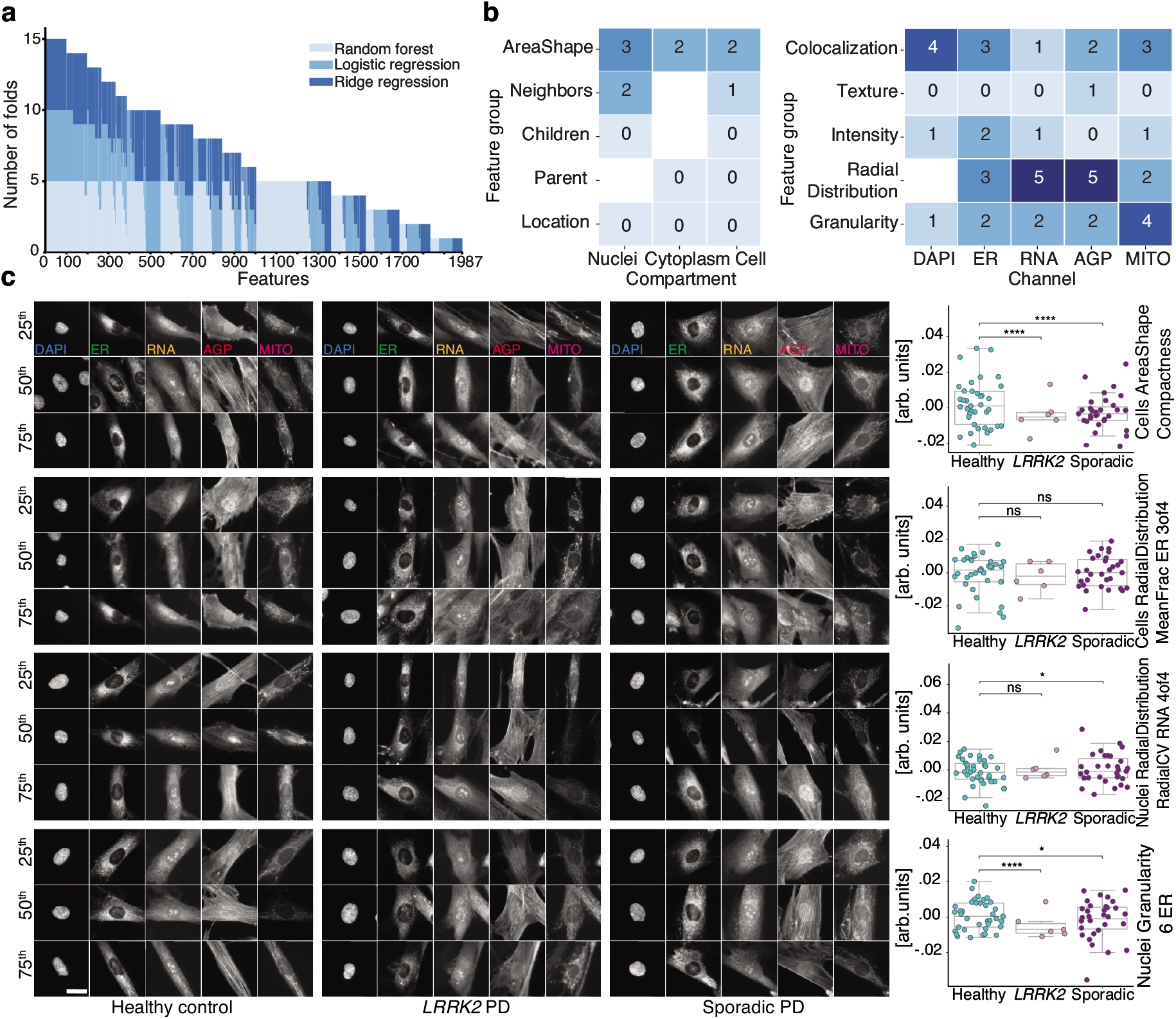
PD classification is driven by a large variety of cell features. **a**, Frequency among 5 crossvalidation folds of 3 models where a CellProfiler feature was within the 1200 most significant of the 3483 features reveals a diverse set of features supporting PD classification. **b**, Frequency of each class of Cell Painting features of the 100 most common features in **a**, with correlated features removed. **c**, Images of representative cells and respective cell line–level mean (n=74 individuals, over all 4 batches) feature values (points and box plot; point colors denote disease state) for 4 features randomly selected from those in **b**. Cells closest to the 25th, 50th and 75th percentiles were selected. Scale bar: 20 μm. Box plot components are: horizontal line, median; box, interquartile range; whiskers, 1.5× interquartile range. arb. units: arbitrary units. Two-sided Mann–Whitney *U* test: ns: *P* > 0.05; *: 0.01 < *P* ≤ 0.05; **”: *P* ≤ 0.0001; *P* values from top to bottom: *P* < 0.0001, *P* < 0.0001, *P* = 0.062, *P* = 0.44, *P* = 0.15, *P* = 0.011, *P* < 0.0001, *P* = 0.012.

From this pool of 55 features, we randomly selected 4 features and inspected their visual and statistical attributes for control, sporadic PD, and *LRRK2* PD cell lines (**Fig. 6c**). Although most of the 55 features were significantly different between control and both *LRRK2* PD (42 had *P* < 0.05, two-sided Mann–Whitney *U* test) and sporadic PD lines (47 had *P* < 0.05, two-sided Mann–Whitney *U* test), there was still considerable variation within each group. Furthermore, these differences were not visually apparent in representative cell images (**Fig. 6c**). Collectively, the results show that the power of our models to accurately classify PD relies on a large number and complex combination of different morphological features, rather than a few salient ones.

## Discussion

To overcome the poor clinical trial outcomes for Parkinson’s disease^27,28^, it is clear that drug discovery strategies that use patient-derived cells^29^ and are less reliant on preconceived hypotheses are required. To address this, we developed a robust platform for the unbiased phenotypic analysis of patient-derived cell lines by applying custom cell culture automation procedures, high-content imaging, and cutting-edge deep learning–based image analysis. While high-throughput image-based profiling has been previously performed using up to 62 cell lines^19^, significant variation in parameters such as cell line growth rate highlights the challenges of large-scale manual cell culture. In contrast, our automated platform enabled the simultaneous profiling of 96 primary cell lines with minimal variation in growth rate characteristics (**Supplementary Fig. 1f**). To our knowledge, this is the first successful demonstration in which automated, unbiased deep learning–based phenotypic profiling is able to discriminate between primary cells from PD patients (both sporadic and *LRRK2*) and healthy controls (**Fig. 5b, c, d**). Interestingly, both deep embeddings and CellProfiler features contained strong disease predictive signatures, evident by PD classification performance that may approach the accuracy limit of PD diagnosis itself^30^. The fact that two divergent analysis approaches both succeed in separating PD fibroblasts from healthy controls provides an independent validation of the results and the experimental pipeline that yielded them.

Furthermore, we demonstrate that the generation of high-content imaging data in parallel with predictive machine learning enabled the individual identification of primary cell lines, even when a second biopsy was provided years later, suggesting the presence of morphological features that are inherent to specific individuals (**Fig. 4c**). This separation between individuals highlights the power of machine learning pattern recognition that is challenging, if not impossible, to capture by human observation alone. Such insights pave the way to uncover novel individual or patient-specific cellular phenotypes, with valuable implications for personalized medicine^31^.

To fortify confidence in our predictions, we applied numerous considerations to reduce noise and confounds^32^. First, we developed a near real-time image quality visualization tool, which has been made publicly available (see Code Availability statement) to maintain consistency across batches and ensure high quality data as input for model generation. Second, all samples were collected by the same organization in a standardized manner, in which all lines underwent equal and rigorous quality control. Third, the experimental design controlled for biases including age, sex, ancestry, passage, biopsy collection, and expansion. We note that all but 5 PD patients reported taking at least one PD-related pharmacological treatment at time of biopsy (**Supplementary Data 1**), and we cannot rule out any possible effects this may have had on fibroblast phenotypes. We used two plate layout designs to randomize samples and control for edge effects, a known confounder. To obtain spatially and temporally independent train and test datasets for cell line identification tasks, the models were trained on data from one plate layout across three batches, while testing was performed on data from the second layout within the fourth held-out batch. The custom automation procedures developed for this project were crucial for achieving data consistency that allowed for cross batch training and validation. Finally, all disease classification tasks were performed with 5-fold cross-validation, using test cell lines from held-out donors, and were thus not influenced by individual signatures.

Deep learning–based image analysis offers unparalleled classification performance; however, defining the specific morphological features that drive our predictions remains challenging. To gain insights into the mechanistic drivers, we performed two comprehensive and complementary analyses to determine which cellular compartments and classes of features were most important for PD classification. First, we repeated the PD classification task, removing the deep embedding dimensions corresponding to one, or all but one, fluorescent channel, and found the former did not have a profound effect on the classification, irrespective of which channel was removed. This indicates interchannel redundancy in observed disease signatures, which could be explained by a combination of signal “bleed-through” between channels and mechanistic interplay between cellular compartments. Second, we evaluated the distribution and types of CellProfiler features important to various models for PD classification (**Fig. 6**). The advantage of this approach is that predictions can potentially be linked directly to specific morphological features. However, our analysis showed that the classification of healthy and PD states relied on over 1200 features, where even the most common significant features were nearly impossible to discern by eye. Taken together, our analysis indicates that the detected PD-specific morphological signatures are extremely complex, much like its clinical manifestation, and comprehensive disruption studies will be required to delineate the underlying molecular mechanisms.

Looking forward, the work we have presented can be extended in many ways. Firstly, to make this large dataset accessible and to avoid overfitting, we summarized the thousands of cells in each well by averaging to a single embedding or feature vector, but this may not capture the heterogeneity that exists among cells in a well. The classification models we used were further purposely chosen to avoid overfitting, but it is possible that more complex models, such as convolutional neural networks trained on cell images, could perform better. The model used to produce the deep embeddings was trained on non-cell images, and perhaps could be retrained or fine-tuned to include cell image datasets. Application of this method to more disease-relevant cell types such as iPSC–derived neurons would be a powerful next step; while variation in the differentiation process can be a concern^32^, it could be mitigated by automated procedures like those presented here and reported previously^20^. Lastly, we expect this approach to be amenable to other cell types, diseases, and morphological stains, and could perhaps benefit from combination with other multi-omics data or additional clinical patient data. In all future applications, reducing technical variation via automation and a robust experimental design will continue to be essential, and we expect the exploration of further computational methods that are robust to nuisance factors (e.g., experimental batch, cell line donor) to be an important area of focus.

The scale of this unbiased high-content profiling experiment is, to our knowledge, unprecedented: it provides the scientific community with the largest publicly available Cell Painting dataset to date (in terms of pixel count) at 48 terabytes in size, compared with the next largest dataset at 13 terabytes (RxRx19a)^33^. Scalable and reproducible automation procedures enabled the generation of high-quality data suited to cross-batch training and validation, an important technical feature for large-scale applications such as drug screening^1^. Our ability to identify Parkinson’s-specific disease signatures using standard cell labeling and deep learning–based image analysis highlights the generalizable potential of this platform to identify complex disease phenotypes in a broad variety of cell types. This represents a powerful, unbiased approach that may facilitate the discovery of precision drug candidates undetectable with traditional target- and hypothesis-driven methods.

## Code availability

Code for generating a deep embedding from an image as well as fitting and evaluating the cell line and PD classification models is available at https://nyscf.org/nyscf-adpd/^39^.

The near real-time image quality analysis is available as a Fiji (an ImageJ distribution) macro at https://github.com/google/microscopeimagequality/tree/main/wellmontagefijimacro^40^.

## Materials availability

Cell lines are available through the NYSCF Repository and can be requested by emailing.

## Data availability

The raw and pre-processed full-resolution images are available upon request due to dataset size constraints, access can be obtained by a data request form at https://nyscf.org/nyscf-adpd/. The processed image, deep embedding, and CellProfiler data are available under the CC BY-NC-SA 4.0 license at https://nyscf.org/nyscf-adpd/.

## Acknowledgments

We thank Geoff Davis and Joshua Cutts for input on analysis; Michael Ando, Patrick Riley, Vikram Khurana, and Phillip Jess for advice on the manuscript; Austin Blanco for help with imaging optimization; Gist Croft for input on sample selection; Yosif Ganat for assistance with cell seeding during platform development; Steve Finkbeiner and Lee Rubin and their teams for helpful discussions from previous collaborations. We are grateful to all of the study participants who donated samples for this research. We dedicate this work to the memory of our dear colleague and friend, Reid Otto (1969–2022).

## Author Contributions

L. S., Y.C., P.C.N., M.F., S.L.S., E.S., S.A.N., F.J.M., Jr., M.B., S.J.Y., and B.J. conceptualized the project. D.C., J.H., E. T., C.J.H., S.D., S.J., P.F., G.B., J.G., R.O., L.A., D.P., and B.J. developed the automation platform. L.S., Y.C., M.F., F. J.M., Jr., D.P., M.B., S.J.Y., and B.J. designed experiments. B.M., D.C., J.H., A.D.L., K.R., and N.G.S.C.A.T. executed the experiments. B.M., C.B., and M.F. developed the image quality analysis software. B.M., Y.C., and C.B. preprocessed image data. B.M., Y.C., M.F., S.A., B.F., A.G., S.V., A.N., Z.A., B.W., M.C., E.A.B., O.P., A.D., M.B., S.J.Y., and B.J. developed models and conducted data analysis. L.S., C.J.H., J.K., L.B., E.S., S.A.N., F.J.M., Jr., D.P., M.B., S.J.Y., and B.J. managed the project. L.S., F.J.M., Jr., D.P., M.B., S.J.Y., and B.J. supervised the work. J.K., S.A.N., F.J.M., Jr., D.P., and M.B. acquired funding and provided resources. L.S., B.M., Z.A., R.S.A., and B.J. created visualizations. L.S., B.M., R.S.A., S.J.Y., and B.J. wrote and edited the manuscript. These authors contributed equally: M. B., S.J.Y., and B.J.

**NYSCF Global Stem Cell Array^®^ Team**

Jenna Hall^2^, Brodie Fischbacher^2^, Christopher J. Hunter^2^, Sean DesMarteau^2^, Selwyn Jacob^2^, Peter Ferrarotto^2^,Geoff Buckley-Herd^2^, Jordan Goldberg^2^, Reid Otto^2^, Alyssa Duren-Lubanski^2^, Kathryn Reggio^2^, Lauren Bauer^2^, Daniel Paull^2^

A full list of members and their affiliations appears in the Supplementary Information.

## Methods

### Donor recruitment and biopsy collection

This project used fibroblasts collected under an IRB-approved protocol at the New York Stem Cell Foundation Research Institute (NYSCF), in compliance with all relevant ethical regulations. After providing written informed consent, participants received a 2–3 mm punch biopsy under local anesthesia performed by a dermatologist at a collaborating clinic. A $50 gift card was provided if the consent version used at the time of collection offered compensation. The dermatologists utilized clinical judgment to determine the appropriate location for the biopsy, with the upper arm being most common. Individuals with a history of scarring and bleeding disorders were ineligible to participate. In addition to biological sample collection, all participants completed a health information questionnaire detailing their personal and familial health history, accompanied by demographic information. All participants with PD self-reported this diagnosis and all but three participants with PD had research records from the same academic medical center in New York available which confirmed a clinical PD diagnosis. To protect participant confidentiality, the biological sample and data were coded and the key to the code securely maintained.

### Experimental design and validation

Cell lines were selected from the NYSCF fibroblast repository containing cell lines from over 1000 participants. We applied strict exclusion criteria based on secondary (non-PD) pathologies, including skin cancer, stroke, epilepsy, seizures, and neurological disorders and, for sporadic PD cases, UPDRS scores below 15. Out of the remaining cell lines, 120 healthy control and PD cell lines were preliminarily matched based on donor age and sex; all donors were self-reported white and most were confirmed to have at least 88% European ancestry via genotyping (**Supplementary Data 1**). The 120 cell lines were all expanded in groups of eight, comprising two pairs of PD and preliminary matched healthy controls, and after expansion was completed, a final set of 96 cell lines, including a set of 45 PD and final matched healthy controls, was selected for the study.

We expanded and froze cells to conduct four identical batches, each consisting of twelve 96-well plates in two unique plate layouts, of which each plate contained exactly one cell line per well. The plate layout consisted of a checkerboard-like pattern of placement of healthy control and Parkinson’s cell lines and cell lines on the edge of the plate in one plate layout were near the center in the other layout (**Fig. 2b**). We populated the plate layouts by randomly permuting the order of the 45 cell line pairs.

To ensure and confirm a balanced plate layout and experimental design, we performed a lasso variable selection for healthy vs. PD in advance of beginning the first experiment batch, to identify covariates that might be good predictors of disease state. Plate layout designs from three random reorderings of the cell line pairs were considered, and the best performing design was selected. Specifically, we sought a design that minimized the covariate weights of a crossvalidated linear regression model with L1 regularization with the following covariates as features: participant age (above or at/below 64 years), sex (male or female), biopsy location (arm, leg, not arm or leg, left, right, not left or right, unspecified), biopsy collection year (at/before or after 2013), expansion thaw freeze date (on/before or after July 11, 2019), thaw format, doubling time (at/less than or greater than 3.07 days), and plate location (well positions not in the center in both layouts, well positions on the edge in at least one plate layout, well positions on a corner in at least one plate layout, row (A/B, C/D, G/E, F/H), column (1–3, 4–6, 7–9, 10–12).

After the experiment was conducted, to further confirm the total number of cells or the growth rates did not represent a potential confound, we reviewed the count of cells, extracted from the CellProfiler analysis, and the doubling time of each cell line by disease state (healthy, sporadic PD, *LRRK2* PD and *GBA* PD). A two-sided Mann–Whitney *U* test, Bonferroni adjusted for 3 comparisons, did not highlight statistical differences.

### Cell line expansion

Each skin biopsy was washed in biopsy plating media containing Knockout-DMEM (Life Technologies #10829-018), 10% FBS (Life Technologies, #100821-147), 2 mM GlutaMAX (Life Technologies, #35050-061), 0.1 mM MEM Non-Essential Amino Acids (Life Technologies, #11140-050), 1X Antibiotic-Antimycotic, 0.1 mM 2-Mercaptoethanol (Life Technologies, #21985-023) and 1% Nucleosides (Millipore, #ES-008-D), dissected into small pieces and allowed to attach to a 6-well tissue culture plate, and grown out for 10 days before being enzymatically dissociated using TrypLE CTS (Life Technologies, #A12859-01) and replated at a 1:1 ratio. Cell density was monitored with daily automated bright-field imaging and upon gaining confluence, cells were harvested and frozen down into repository vials at a density of 100,000 cells per vial in 1.5 mL of CTS Synth-a-Freeze (Life Technologies, #A13717-01) in an automated fashion using the NYSCF Global Stem Cell Array^® 20^.

To expand cells for profiling, custom automation procedures were developed on an automation platform consisting of a liquid handling system (Hamilton STAR) connected to a Cytomat C24 incubator, a Celigo cell imager (Nexcelom), a VSpin centrifuge (Agilent), and a Matrix tube decapper (Hamilton Storage Technologies). Repository vials were thawed manually in two batches of 4, for a total of 8 lines per run. To reduce the chance of processing confounds, when possible, every other line that was processed was a healthy control, the order of lines processed alternated between expansion groups, and the scientist performing the expansion was blinded to the experimental group. Repository tubes were placed in a 37 °C water bath for 1 minute. Upon removal, fibroblasts were transferred to their respective 15 mL conical tubes at a 1:2 ratio of Synth-a-Freeze and Fibroblast Expansion Media (FEM). All 8 tubes were spun at 1100 RPM for 4 minutes. Supernatant was aspirated and resuspended in 1 mL FEM for cell counting, whereby an aliquot of the supernatant was incubated with Hoechst (H3570, ThermoFisher) and Propidium Iodide (P3566, ThermoFisher) before being counted using a Celigo automated cell imager. Cells were plated in one well of a 6-well at 85,000–120,000 cells in 2 mL of FEM. If the count was lower than 75,000, cells were plated into a 12-well plate and given the appropriate amount of time to reach confluence. Upon reaching 90-100% confluence, the cell line was added into another group of 8 to enter the automated platform. All 6-well and 12-well plates were kept in a Cytomat C24 incubator and every passage and feed from this point onward was automated (Hamilton STAR). Each plate had a FEM media exchange every other day and underwent passages every 7th day. The cells were fed with FEM using an automated method that retrieved the plates from the Cytomat two at a time and exchanged the media.

After 7 days, the batch of 8 plates had a portion of their supernatant removed and banked for mycoplasma testing. Cells were passaged and plated at 50,000 cells per well (into up to 6 wells of a 6 well plate) and allowed to grow for another 7 days. Not every cell line was expected to reach the target of filling an entire 6-well plate. To account for this, a second passage at a fixed seeding density of 50,000 cells per well was embedded in the workflow for all of the lines. After another 7 days, each line had a full 6-well plate of fibroblasts and generated a minimum of 5 assay vials with 100,000 cells per vial. The average doubling time for each cell line was calculated by taking the log base 2 of the ratio of the cell number at harvest over the initial cell number. Each line was then propagated a further two passages and harvested to cryovials for DNA extraction.

### Automated screening

Custom automation procedures were developed for large-scale phenotypic profiling of primary fibroblasts. For each of the four experimental batches, 2D barcoded matrix vials from 96 lines containing 100,000 cells per vial were thawed, decapped and rinsed with FEM. Cells were spun down at 192 g for 5 minutes, supernatant was discarded, and cells were resuspended in culture media. Using a Hamilton Star liquid handling system, the cells were then seeded onto five 96-well plates (Fisher Scientific, 07-200-91) for post-thaw recovery. Cells were harvested 5 days later using automated methods as previously described^20^. In brief, media was removed from the cells and rinsed with TrypLE. Cells were incubated for 30 minutes at 37 before being neutralized with FEM. Cells were centrifuged before supernatants were aspirated and cells resuspended in FEM. An aliquot of cell suspension was incubated with Hoechst (H3570, ThermoFisher) and Propidium Iodide (P3566, ThermoFisher) before being counted using a Celigo automated imager. Using an automated seeding method developed on a Lynx liquid handling system (Dynamic Devices, LMI800), cell counts from each line were used to adjust cell densities across all 96 lines to transfer a fixed amount of cells into two 96-well deep well troughs in two distinct plate layouts. Each layout was then stamped onto six 96-well imaging plates (CellVis, P96-1.5H-N) at a fixed target density of 3,000 cells per well. Assay plates were then transferred to a Cytomat C24 incubator for two days before phenotypic profiling where cells were stained and imaged as described below. All cell lines were screened at a final passage number of 10 or 11 +/− 2. In total, this process took 7 days and could be executed by a single operator.

### Staining and imaging

To fluorescently label the cells, the protocol published in Bray et al.^21^ was adapted to an automated liquid handling system (Hamilton STAR). Briefly, plates were placed on deck for addition of culture medium containing MitoTracker (Invitrogen^™^ M22426) and incubated at 37 °C for 30 minutes, then cells were fixed with 4% Paraformaldehyde (Electron Microscopy Sciences, 15710-S), followed by permeabilization with 0.1% Triton X-100 (Sigma-Aldrich, T8787) in 1X HBSS (Thermo Fisher Scientific, 14025126). After a series of washes, cells were stained at room temperature with the Cell Painting staining cocktail for 30 minutes, which contains Concanavalin A, Alexa Fluor^®^ 488 Conjugate (Invitrogen^™^ C11252), SYTO^®^ 14 Green Fluorescent Nucleic Acid Stain (Invitrogen^™^ S7576), Alexa Fluor^®^ 568 Phalloidin (Invitrogen^™^ A12380), Hoechst 33342 trihydrochloride, trihydrate (Invitrogen^™^ H3570), Molecular Probes Wheat Germ Agglutinin, Alexa Fluor 555 Conjugate (Invitrogen^™^ W32464). Plates were washed twice and imaged immediately.

The images were acquired using an automated epifluorescence system (Nikon Ti2). For each of the 96 wells acquired per plate, the system performed an autofocus task in the ER channel, which provided dense texture for contrast, in the center of the well, and then acquired 76 non-overlapping tiles per well at a 40× magnification (Olympus CFI-60 Plan Apochromat Lambda 0.95 NA). To capture the entire Cell Painting panel, we used 5 different combinations of excitation illumination (SPECTRA X from Lumencor) and emission filters (395 nm and 447/60 nm for Hoechst, 470 nm and 520/28 nm for Concanavalin A, 508 nm and 593/40 nm for RNA-SYTO14, 555 nm and 640/40 nm for Phalloidin and wheat-germ agglutinin, and 640 nm and 692/40 nm for MitoTracker Deep Red). Each 16-bit 5056×2960 tile image was acquired using NIS-Elements AR acquisition software from the image sensor (Photometrics Iris 15, 4.25 μm pixel size). Each 96-well plate resulted in approximately 1 terabyte of data.

### Near real-time image quality analysis

To assess the quality and consistency of the images collected from a full 96-well plate, we developed a near real-time Fiji (an ImageJ distribution) macro^34^. The tool creates image montages from random image crops from each channel across all wells on a plate with related focus scores and intensity statistics. These montages were inspected to confirm that images were suitable as input for further analysis. The tool is able to generate a representative sampling of images from one 96-well plate (1 terabyte) in 30 minutes.

### Confirming cell line provenance

All 96 lines were analyzed using NeuroChip^35^ or similar genome-wide SNP genotyping arrays to check for PD-associated mutations (*LRRK2* G2019S and *GBA* N370S). PD lines that did not contain *LRRK2* or *GBA* mutations were classified as Sporadic. NeuroChip analysis confirmed the respective mutations for all lines from *LRRK2* and *GBA* PD individuals, with the exceptions of cell line 48 from donor 10124, where no *GBA* mutation was detected, and the control cell line 77 (from donor 51274) where an N370S mutation was identified. This prompted a post hoc ID SNP analysis (using Fluidigm SNPTrace) of all expanded study materials, which confirmed the lines matched the original ID SNP analysis made at the time of biopsy collection for all but two cell lines: cell line 48 from donor 10124 (*GBA* PD) and cell line 57 from donor 50634 (healthy), which have been annotated as having unconfirmed cell line identity in **Supplementary Data 1**. We confirmed the omission of line 48 and 77 did not qualitatively impact *GBA* PD vs healthy classification (**Supplementary Figure 4**) and although line 57 was most likely from another healthy individual, we confirmed the omission of line 57 had minimal impact, yielding a 0.77 (0.08 SD) ROC AUC (compared with 0.79 (0.08 SD) from including the line) for *LRRK2*/Sporadic PD vs. healthy classification (logistic regression trained on tile deep embeddings). Importantly, the post hoc ID SNP analysis did confirm the uniqueness of all 96 lines in the study. Finally, for a subset of 89 of the 96 lines, which were genotyped using the NeuroChip, we additionally confirmed none of these lines contained any other variants reported in ClinVar to have a causal, pathogenic association with PD, across mutations spanning genes *GBA*, *LRRK2*, *MAPT*, *PINK1*, *PRKN* and *SNCA* (except those already reported to carry G2019S (*LRRK2*) and N370S (*GBA*)).

### Image statistics features

For assessing data quality and baseline predictive performance on classification tasks, we computed various image statistics. Statistics are computed independently for each of the 5 channels for the image crops centered on detected cell objects. For each tile or cell, a “focus score” between 0.0 and 1.0 was assigned using a pre-trained deep neural network model^36^. Otsu’s method was used to segment the foreground pixels from the background and the mean and standard deviation of both the foreground and background were calculated. Foreground fraction was calculated as the number of foreground pixels divided by the total pixels. All features were normalized by subtracting the mean of each batch and plate layout from each feature and then scaling each feature to have unit L2 norm across all examples.

### Image pre-processing

We first flat field–corrected 16-bit images by obtaining an estimate of the background intensity by taking the 10th percentile image across all images from the same batch, plate and channel, blurred with a Gaussian kernel of sigma 50, and then dividing each image by this background intensity estimate. Next, Otsu’s method was used in the DAPI channel to detect nuclei centers. Images were converted to 8-bit after clipping at the 0.001 and 1.0 minimum and maximum percentile values per channel and applying a log transformation. These 8-bit 5056×2960×5 images, along with 512×512×5 image crops centered on the detected nuclei, were used to compute deep embeddings. Only image crops existing entirely within the original image boundary were included for deep embedding generation.

### Deep image embedding generation

Deep image embeddings were computed on both the tile images and the 512×512×5 cell image crops. In each case, for each image and each channel independently, we first duplicated the single channel image across the RGB (red-green-blue) channels and then inputted the 512×512×3 image into an Inception architecture^22^ convolutional neural network, pre-trained on the ImageNet^22,23^ object recognition dataset consisting of 1.2 million images of a thousand categories of (non-cell) objects, and then extracted the activations from the penultimate fully connected layer and took a random projection to get a 64-dimensional deep embedding vector (64×1×1). We concatenated the five vectors from the 5 image channels to yield a 320-dimensional vector or embedding for each tile or cell crop. 0.7% of tiles were omitted because they were either in wells never plated with cells due to shortages or because no cells were detected, yielding a final dataset consisting of 347,821 tile deep embeddings and 5,813,995 cell image deep embeddings. All deep embeddings were normalized by subtracting the mean of each batch and plate layout from each deep embedding. Finally, we computed datasets of the well-mean deep embeddings, the mean across all cell or tile deep embeddings in a well, for all wells.

### CellProfiler feature generation

We used a CellProfiler pipeline template^21^ where we determined Cells in the RNA channel, Nuclei in the DAPI channel and Cytoplasm by subtracting the Nuclei objects from the Cell objects. We ran CellProfiler^24^ version 3.1.5^37^ independently on each 16-bit 5056×2960×5 tile image set, inside a Docker container on Google Cloud. 0.2% of the tiles resulted in errors after multiple attempts and were omitted. Features were concatenated across Cells, Cytoplasm and Nuclei to obtain a 3483-dimensional feature vector per cell, across 7,450,738 cells. As we were unable to load this dataset and fit a model^38^ in memory with 196 gigabytes of memory, we computed a reduced dataset with the well-mean feature vector per well. We then normalized all features by subtracting the mean of each batch and plate layout from each feature and then scaled each feature to have unit L2 norm across all examples.

### Modeling and analysis

We evaluated several classification tasks ranging from cell line prediction to disease state prediction using various data sources and multiple classification models. Data sources consisted of image statistics, CellProfiler features and deep image embeddings. Since data sources and predictions could have existed at different levels of aggregation ranging from the cell–level, tile-level, well-level to cell line–level, we used well-mean aggregated data sources (averaging all cell features or tile embeddings in a well) as input to all classification models, and aggregated the model predictions by averaging predicted probability distributions (the cell line–level prediction, by averaging predictions across wells for a cell line). In each classification task, we defined an appropriate cross-validation approach and all figures of merit reported are those on the held-out test sets. For example, the well-level accuracy is the accuracy of the set of model predictions on the held out wells, and the cell line–level accuracy is the accuracy of the set of cell line–level predictions from held out wells. The former indicates the expected performance with just one well example, while the latter indicates expected performance from averaging predictions across multiple wells; any gap could be due to intrinsic biological, process or modeling noise and variation.

Various classification models (sklearn) were used, including a cross-validated logistic regression (solver = “lbfgs”, max_iter = 1000000), random forest classifier (with 100 base estimators), cross-validated ridge regression and multilayer perceptron (single hidden layer with 200 neurons, max_iter = 1000000); these settings ensured solver convergence to the default tolerance.

### Cell line identification analysis

For each of the various data sources, we utilized the crossvalidation sets defined in **Supplementary Table 1**. For each train/test split, one of several classification models was fit or trained to predict a probability distribution across the 96 classes, the ID of the 96 unique cell lines. For each prediction, we evaluated both the top predicted cell line, the cell line class to which the model assigns highest probability, as well as the predicted rank, the rank of probability assigned to the true cell line (when the top predicted cell line is the correct one, the predicted rank is 1). We used as the figure of merit the well-level or cell line–level accuracy, the fraction of wells or cell lines for which the top predicted cell line among the 96 possible choices was correct.

### Biopsy donor identification analysis

For each of the various data sources, we utilized the cross-validation sets defined in **Supplementary Table 2**. For each train/test split, one of several classification models was fit or trained to predict a probability distribution across 91 classes, the possible donors from which a given cell line was obtained. For each of the 5 held-out cell lines, we evaluated the cell line–level predicted rank (the predicted rank assigned to the true donor).

### *LRRK2* and sporadic PD classification analysis

For each of the various data sources, we partitioned the demographically-matched healthy/PD cell line pairs into 5 groups with a near-even distribution of PD mutation, sex and age, which were then used as folds for cross-validation (**Supplementary Data 2**). For a given group, we trained a model on the other 4 groups on a binary classification task, healthy vs. PD, before testing the model on the held-out group of cell line pairs. The model predictions on the held-out group were used to compute a receiver operator characteristic (ROC) curve, for which the area under the curve (ROC AUC) can be evaluated. The ROC curve is the true positive rate vs. false positive rate, evaluated at different predicted probability thresholds. ROC AUC can be interpreted as the probability of correctly ranking a random healthy control and PD cell line. The ROC AUC was computed for cell line–level predictions, the average of the models’ predictions for each well from each cell line. We evaluated the ROC AUC for a given held-out fold in three ways: with model predictions for both all sporadic and *LRRK2* PD vs. all controls, all *LRRK2* PD vs. all controls, and all sporadic PD vs. all controls. We obtained overall ROC AUC by taking the average and standard deviation across the 5 cross-validation sets.

### PD classification analysis with *GBA* PD cell lines

For a preliminary analysis (**Supplementary Figure 4**) only, the PD vs. healthy classification task was conducted with a simplified crossvalidation strategy, where matched PD and healthy cell line pairs were randomly divided into a train half and a test half 8 times. This was done for all matched cell line pairs, just *GBA* PD and matched controls, just *LRRK2* PD and matched controls, and just sporadic PD and matched controls. Test set ROC AUC was evaluated as in the above analysis.

### CellProfiler feature importance analysis

First, we estimated the threshold for number of top-ranked CellProfiler features for a random forest classifier (1000 base estimators) required to maintain the same classification performance as the full set of 3483 CellProfiler features, by evaluating performance for sets of features increasing in size in increments of 20 features.

After selecting 1200 as the threshold, we looked at the top 1200 features for each of the logistic regression, ridge regression and a random forest classifier models. The 100 CellProfiler features shared in common across all five folds of all three model architectures were further filtered using a Pearson’s correlation value threshold of 0.75, leaving 55 features and subsequently grouped based on semantic properties. A feature was selected at random from each of 4 randomly selected groups to inspect the distribution of their values and representative cells from each disease state, with the closest value to the distribution median and quantiles, were selected for inspection. The statistical differences were evaluated using a two-sided Mann–Whitney *U* test, Bonferroni adjusted for 2 comparisons.

