## Supplementary tables and figures for "Integrating deep learning and unbiased automated high-content screening to identify complex disease signatures in human fibroblasts"

---

[1]Google Research, Mountain View, California, USA. [2]The New York Stem Cell Foundation Research Institute, New York, New York, USA. [3]These authors contributed equally: Lauren Schiff, Bianca Migliori, Ye Chen, Deidre Carter. *A list of authors and their affiliations appears at the end of the paper.

NYSCF Global Stem Cell Array Team members: Barry McCarthy[2], Camille Fulmore[2], Daria LeGall[2], Brandon Pearl[2], Dong Woo Shin[2], Dorota Moroziewicz[2], Tomasz Rusielewicz[2], Patrick Fenton[2], Paul McCoy[2], Jenna Hall[2], Brodie Fischbacher[2], Christopher J. Hunter[2], Sean DesMarteau[2], Selwyn Jacob[2], Peter Ferrarotto[2], Geoff Buckley-Herd[2], Reid Otto[2], Jordan Goldberg[2], Kathryn Reggio[2], Alyssa Duren-Lubanski[2], Lauren Bauer[2], Daniel Paull[2]

| | | | Cross-validation | | | | | | | |
|---|---|---|---|---|---|---|---|---|---|---|
| | | | set #1 | set #2 | set #3 | set #4 | set #5 | set #6 | set #7 | set #8 |
| Batch | Plate layout | Cell lines | | | | | | | | |
| | 1 | all 96 | test | ignore | ignore | ignore | ignore | train | train | train |
| 1 | 2 | all 96 | ignore | train | train | train | test | ignore | ignore | ignore |
| | 1 | all 96 | ignore | test | ignore | ignore | train | ignore | train | train |
| 2 | 2 | all 96 | train | ignore | train | train | ignore | test | ignore | ignore |
| | 1 | all 96 | ignore | ignore | test | ignore | train | train | ignore | train |
| 3 | 2 | all 96 | train | train | ignore | train | ignore | ignore | test | ignore |
| | 1 | all 96 | ignore | ignore | ignore | test | train | train | train | ignore |
| 4 | 2 | all 96 | train | train | train | ignore | ignore | ignore | ignore | test |

**Supplementary Table 1 | Cross-validation strategy for 96-way cell line classification.** For each of 8 cross-validation sets, both batch and plate layout were held out in the test set.

| | | | Cross-validation | | | | | | | |
|---|---|---|---|---|---|---|---|---|---|---|
| | | | set #1 | set #2 | set #3 | set #4 | set #5 | set #6 | set #7 | set #8 |
| Batch | Plate layout | Cell lines | | | | | | | | |
| | | 5 held-out biopsies | test | ignore | ignore | ignore | ignore | ignore | ignore | ignore |
| | 1 | remaining 91 lines | ignore | ignore | ignore | ignore | ignore | train | train | train |
| | | 5 held-out biopsies | ignore | ignore | ignore | ignore | test | ignore | ignore | ignore |
| 1 | 2 | remaining 91 lines | ignore | train | train | train | ignore | ignore | ignore | ignore |
| | | 5 held-out biopsies | ignore | test | ignore | ignore | ignore | ignore | ignore | ignore |
| | 1 | remaining 91 lines | ignore | ignore | ignore | ignore | train | ignore | train | train |
| | | 5 held-out biopsies | ignore | ignore | ignore | ignore | ignore | test | ignore | ignore |
| 2 | 2 | remaining 91 lines | train | ignore | train | train | ignore | ignore | ignore | ignore |
| | | 5 held-out biopsies | ignore | ignore | test | ignore | ignore | ignore | ignore | ignore |
| | 1 | remaining 91 lines | ignore | ignore | ignore | ignore | train | train | ignore | train |
| | | 5 held-out biopsies | ignore | ignore | ignore | ignore | ignore | ignore | test | ignore |
| 3 | 2 | remaining 91 lines | train | train | ignore | train | ignore | ignore | ignore | ignore |
| | | 5 held-out biopsies | ignore | ignore | ignore | test | ignore | ignore | ignore | ignore |
| | 1 | remaining 91 lines | ignore | ignore | ignore | ignore | train | train | train | ignore |
| | | 5 held-out biopsies | ignore | ignore | ignore | ignore | ignore | ignore | ignore | test |
| 4 | 2 | remaining 91 lines | train | train | train | ignore | ignore | ignore | ignore | ignore |

**Supplementary Table 2 | Cross-validation strategy for 91-way biopsy donor classification.** For each of 8 cross-validation sets, the test set consisted of cell lines from one of the two biopsies from the 5 individuals who donated two biopsies, while the train set consisted of cell lines from the complementary set of biopsies from these 5 individuals and the remaining 86 individuals who donated only a single biopsy. To avoid plate position biases as potential confounds, plate layout was also held out, and to assess model generalization to a test biopsy acquired in a new batch, batch was also held out. These 8 cross-validation sets were conducted twice, once holding out in the test sets the earlier set of skin biopsies from the 5 individuals who donated two biopsies (cell lines 08, 39, 51, 55, 70), and again holding out the later set (cell lines 91, 92, 93, 94, 95).

cells_AreaShape_Compactness
cells_AreaShape_Eccentricity
cytoplasm_AreaShape_Compactness
cytoplasm_AreaShape_Eccentricity

cells_AreaShape_Zernike_8_4

cells_Correlation_K_ER_RNA
cytoplasm_Correlation_K_ER_RNA
cytoplasm_Correlation_K_ER_DNA

cells_Correlation_Overlap_DNA_ER

cells_Correlation_Overlap_ER_RNA

cells_Correlation_Overlap_Mito_ER
cytoplasm_Correlation_Overlap_Mito_ER

cells_Correlation_RWC_RNA_Mito

cells_Granularity_6_AGP
cells_Granularity_7_AGP

cells_Intensity_IntegratedIntensity_Mito
cytoplasm_Intensity_MassDisplacement_AGP
cytoplasm_AreaShape_Area
cytoplasm_Intensity_IntegratedIntensity_RNA
cells_AreaShape_MeanRadius
cells_AreaShape_MaximumRadius
cytoplasm_Intensity_IntegratedIntensityEdge_Mito
cells_Intensity_IntegratedIntensityEdge_Mito
cytoplasm_Intensity_IntegratedIntensity_Mito
cells_AreaShape_Area
cytoplasm_Intensity_IntegratedIntensity_DNA
cells_Intensity_IntegratedIntensity_RNA
cytoplasm_Intensity_IntegratedIntensity_ER

cells_Intensity_MassDisplacement_DNA

cells_Neighbors_PercentTouching_5
cells_Neighbors_PercentTouching_Adjacent
cells_Neighbors_NumberOfNeighbors_5
cells_Neighbors_NumberOfNeighbors_Adjacent

cells_RadialDistribution_FracAtD_AGP_1of4

cells_RadialDistribution_MeanFrac_ER_3of4

cells_RadialDistribution_MeanFrac_RNA_4of4

cytoplasm_AreaShape_Solidity
cytoplasm_AreaShape_Extent
cells_AreaShape_Solidity

cytoplasm_AreaShape_Zernike_6_4
cytoplasm_AreaShape_Zernike_8_8
cells_AreaShape_Zernike_6_6
cytoplasm_AreaShape_Zernike_6_6

cytoplasm_Correlation_Correlation_Mito_ER
cells_Correlation_Correlation_Mito_ER

cytoplasm_Correlation_Manders_AGP_DNA
cytoplasm_Correlation_Manders_RNA_DNA
cytoplasm_Correlation_Manders_Mito_DNA

cytoplasm_Correlation_RWC_DNA_AGP

cytoplasm_Granularity_10_RNA

cytoplasm_Granularity_1_AGP
cells_Granularity_1_AGP

cytoplasm_RadialDistribution_MeanFrac_AGP_1of4
cytoplasm_RadialDistribution_MeanFrac_AGP_3of4

cytoplasm_RadialDistribution_MeanFrac_RNA_3of4
cells_Granularity_15_RNA

cytoplasm_RadialDistribution_RadialCV_AGP_3of4

cytoplasm_RadialDistribution_RadialCV_RNA_2of4

nuclei_AreaShape_Zernike_2_0

nuclei_AreaShape_Zernike_4_2

nuclei_AreaShape_Zernike_7_1

nuclei_Correlation_Correlation_DNA_RNA

nuclei_Correlation_Correlation_ER_AGP
nuclei_Correlation_Correlation_Mito_AGP

nuclei_Correlation_Manders_AGP_DNA
nuclei_Correlation_Manders_RNA_DNA

nuclei_Correlation_Manders_Mito_ER
nuclei_Correlation_Manders_RNA_ER

nuclei_Correlation_Overlap_DNA_AGP

nuclei_Granularity_11_ER
nuclei_Granularity_8_ER
nuclei_Granularity_7_ER

nuclei_Granularity_13_Mito

nuclei_Granularity_14_DNA

nuclei_Granularity_2_Mito

nuclei_Granularity_4_Mito

nuclei_Granularity_6_ER

nuclei_Granularity_8_Mito

nuclei_Granularity_8_RNA

nuclei_Intensity_IntegratedIntensityEdge_ER
nuclei_Texture_Contrast_ER_10_02

nuclei_Intensity_IntegratedIntensityEdge_RNA
nuclei_Intensity_IntegratedIntensity_RNA

nuclei_Intensity_IntegratedIntensity_ER

nuclei_Neighbors_NumberOfNeighbors_1

nuclei_Neighbors_SecondClosestDistance_1

nuclei_RadialDistribution_FracAtD_RNA_3of4

nuclei_RadialDistribution_MeanFrac_AGP_2of4

nuclei_RadialDistribution_MeanFrac_Mito_3of4
nuclei_Correlation_Correlation_DNA_Mito

nuclei_RadialDistribution_MeanFrac_Mito_4of4

nuclei_RadialDistribution_RadialCV_AGP_3of4

nuclei_RadialDistribution_RadialCV_ER_1of4

nuclei_RadialDistribution_RadialCV_ER_3of4
nuclei_RadialDistribution_RadialCV_ER_2of4

nuclei_RadialDistribution_RadialCV_RNA_4of4
cells_RadialDistribution_RadialCV_RNA_3of4

nuclei_Texture_Correlation_AGP_10_01
nuclei_Texture_InfoMeas2_AGP_10_01
nuclei_Texture_InfoMeas2_AGP_10_03

**Supplementary Table 3 | Most common important CellProfiler features grouped based on correlation.** The top 100 most important CellProfiler features from **Fig. 6a**, filtered down to 55, based on Pearson correlation, and then grouped semantically.

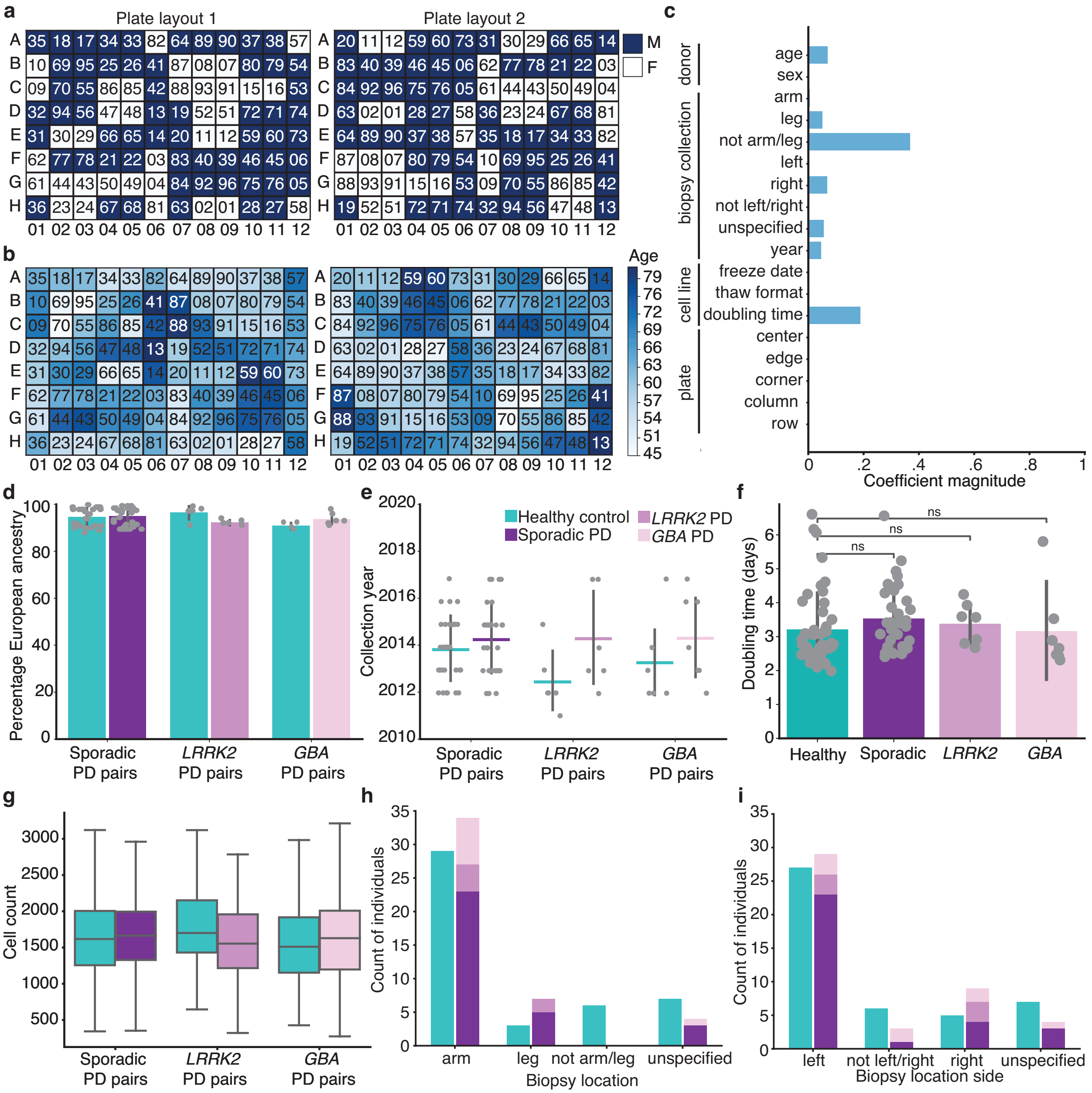


**Supplementary Fig. 1 | Experiment design details for high-content screening.** Various donor demographics including (**a**) sex (male (M), female (F)) and (**b**) age for the two 96-well plate layouts, where each well contains cells from the cell line denoted by the two-digit label. **c**, Lasso variable selection for healthy vs. PD on donor, biopsy, cell line, and plate covariates reveals no significant biases. Distributions of additional cell line covariates including (**d**) percentage European ancestry from genotyping analysis, (**e**) biopsy collection year, (**f**) cell doubling times (two-sided Mann–Whitney $U = 57.0$, $P = 0.01$ for sporadic, $U = 118.0$, $P = 0.64$ for *LRRK2 PD*, and $U = 193.5$, $P = 1.00$ for *GBA PD* vs. healthy, respectively, ns: $P > 0.05$), (**g**) well-level cell count, and biopsy location, (**h**) arm or leg and (**i**) left or right. Box plot components are: horizontal line, median; box, interquartile range; whiskers, 1.5× interquartile range. Bar plot data are presented as mean values +/- standard deviation.

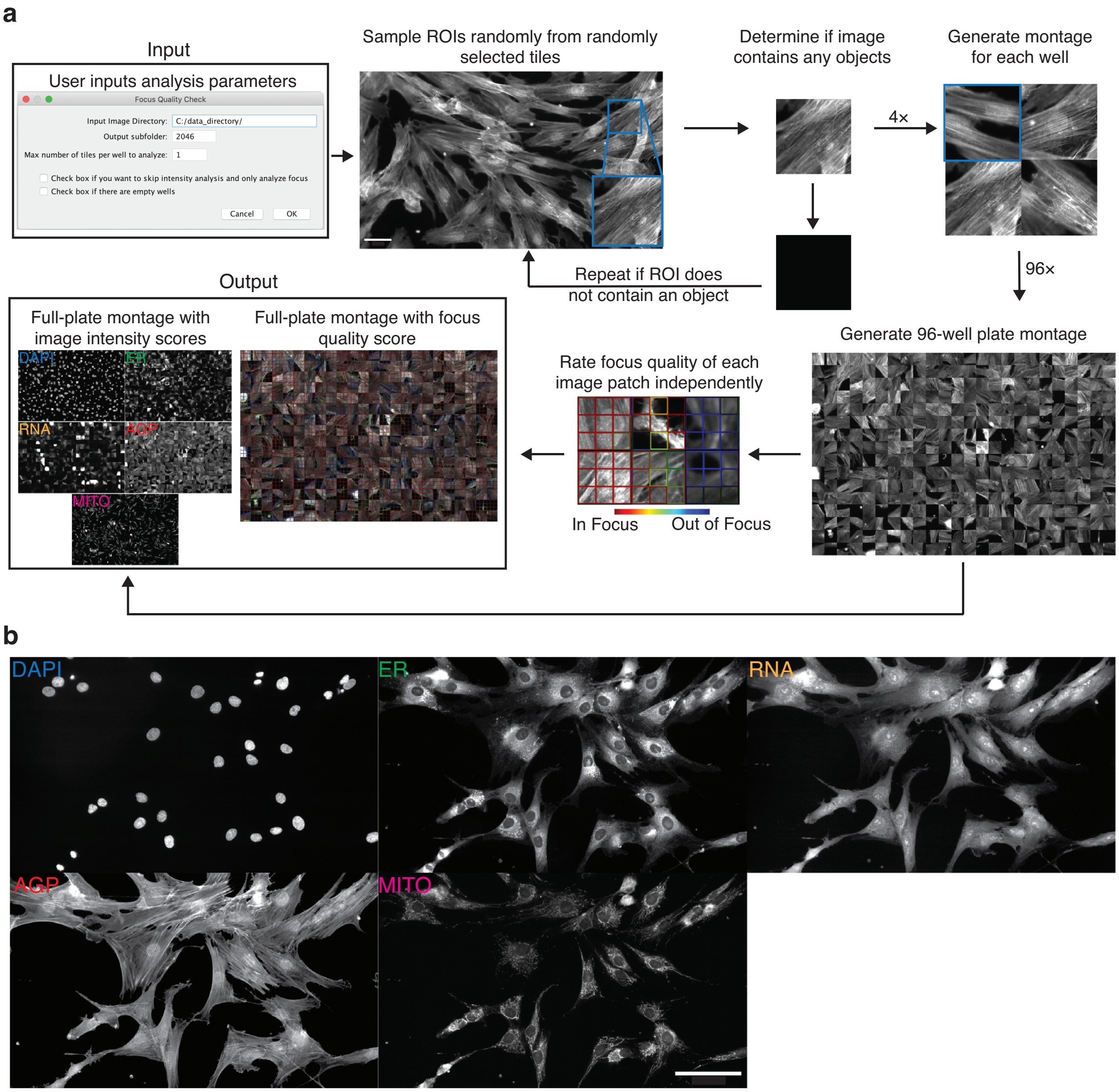


**Supplementary Fig. 2 | Overview of near real-time image quality analysis and sample Cell Painting images of primary human fibroblasts.** A Fiji (an ImageJ distribution) macro assesses the quality and consistency of the images sampled from a full 96-well plate. **a**, Four random regions of interest (ROI) are cropped from images in each channel and in each well, and 96-well montages are constructed for viewing. A measurement of mean image intensity across the plate is reported for each plate montage. Next, the montage corresponding to the user-designated focus channel is inputted to a microscope image focus classifier which calculates a focus quality score for each image patch. For visualization, a color-coded overlay on top of the montage highlights regions that are in focus (red) or out of focus (blue). Scale bar: 50 µm. **b**, Sample images of one tile from the 5 Cell Painting channels. Sample images are representative of those from 48 experimental well plates across 4 experimental batches. Scale bar: 100 µm.

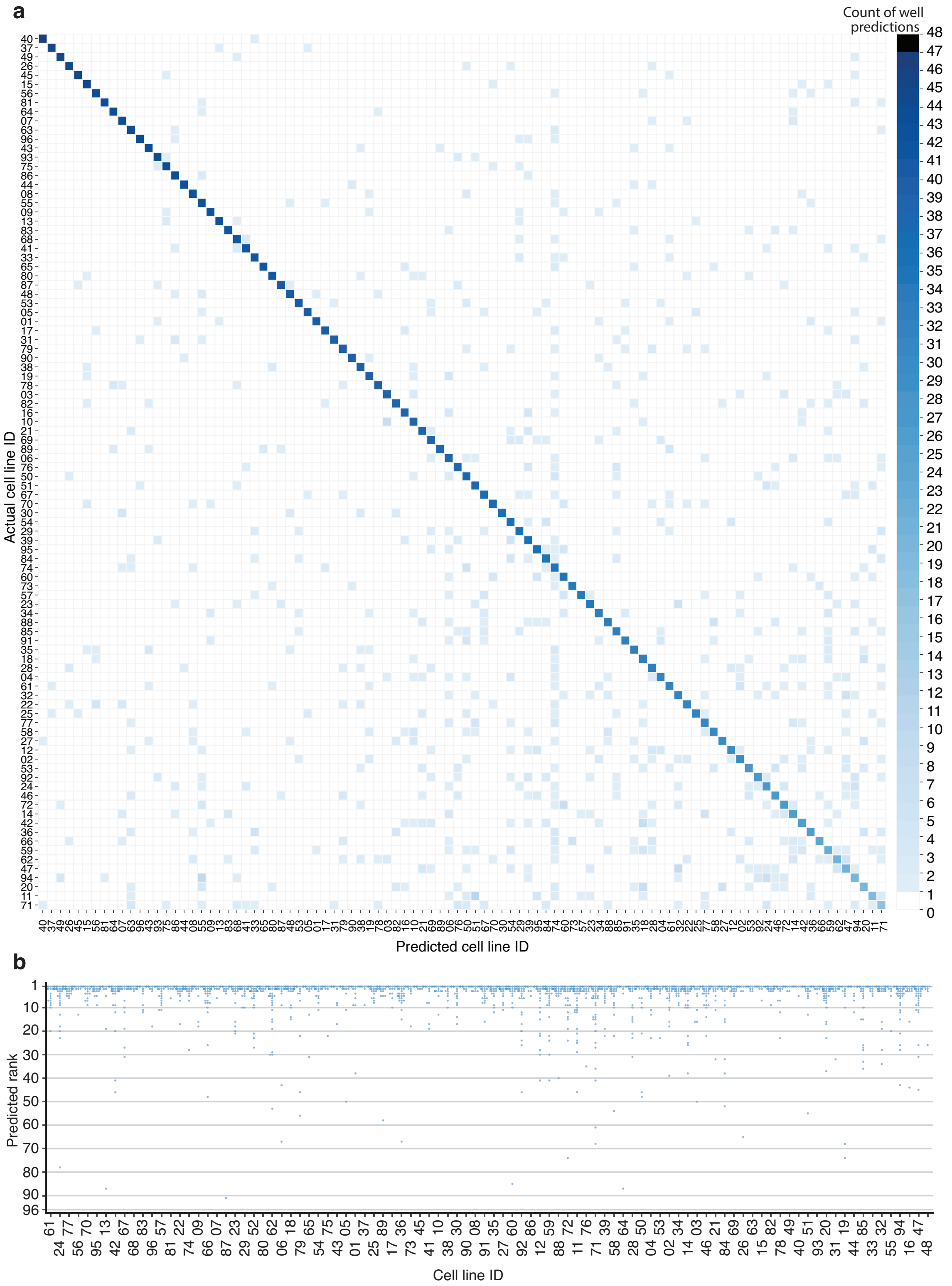


**Supplementary Fig. 3 | Identification of individual cell lines in held-out batches and plate layouts at the well-level. a**, Confusion matrix, sorted by the diagonal, showing the test set well-level predicted and actual cell lines for each of 6 wells in each of 8 held-out batch and held-out plate layouts for the model in **Fig. 3c**. **b**, Test set well-level predicted rank, among 96 of the 6 wells in each of 8 held-out batch and held-out plate layouts for the model in **Fig. 3c**.

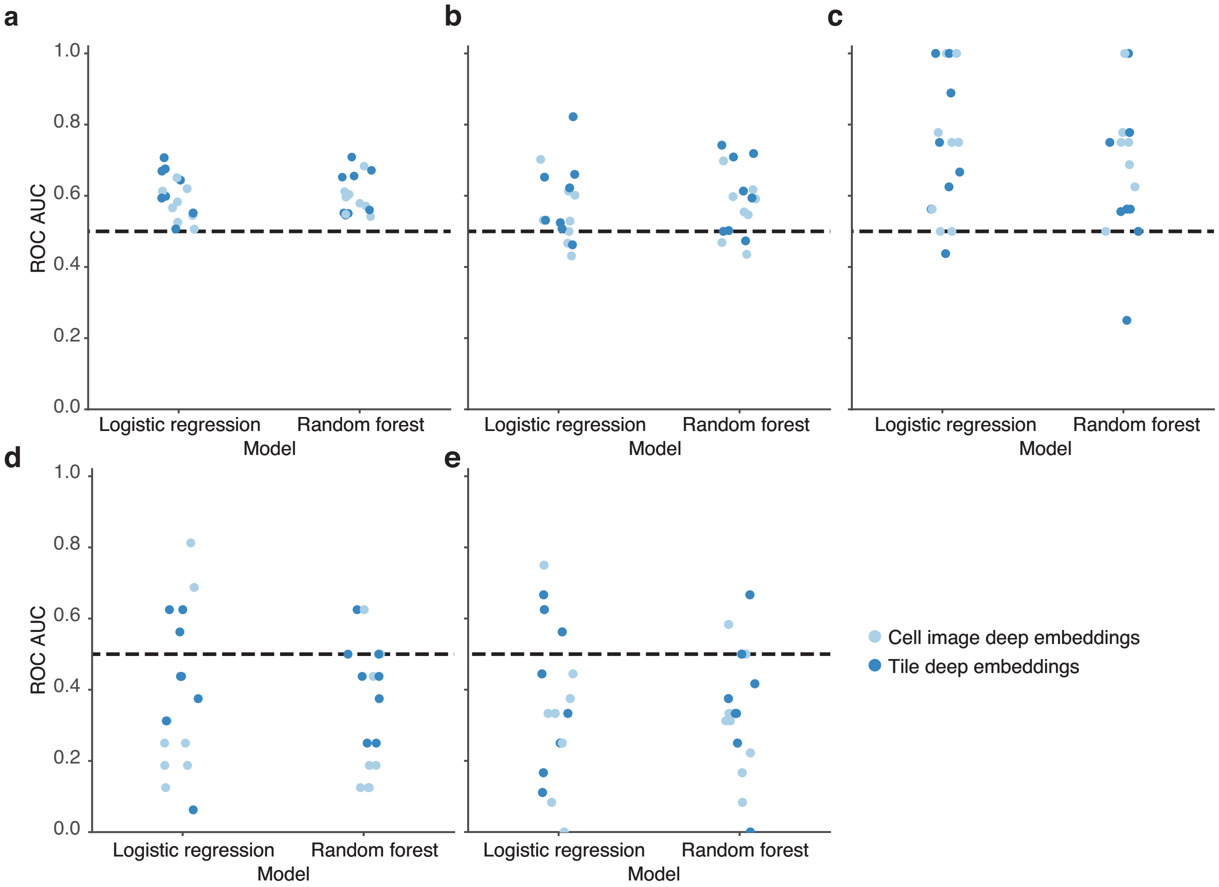


**Supplementary Fig. 4 | Preliminary evaluation of PD classification performance.** Test set cell line–level PD classification for (**a**) all PD (*n* = 45 participants) and matched controls (n = 45 participants), (**b**) sporadic PD (*n* = 31) and matched controls (*n* = 31 participants), (**c**) *LRRK2* PD (*n* = 6 participants) and matched controls (*n* = 6 participants), (**d**) *GBA* PD (*n* = 8 participants) and matched controls (*n* = 8 participants), and (**e**) *GBA* PD (*n* = 7 participants) and matched controls (*n* = 7 participants), excluding cell lines 48 (unconfirmed *GBA*) and 77 (Healthy with *GBA*); see Methods. In each case, for cross-validation, matched cell line pairs were randomly divided into a train half and a test half 8 times. Dashed line denotes chance performance.

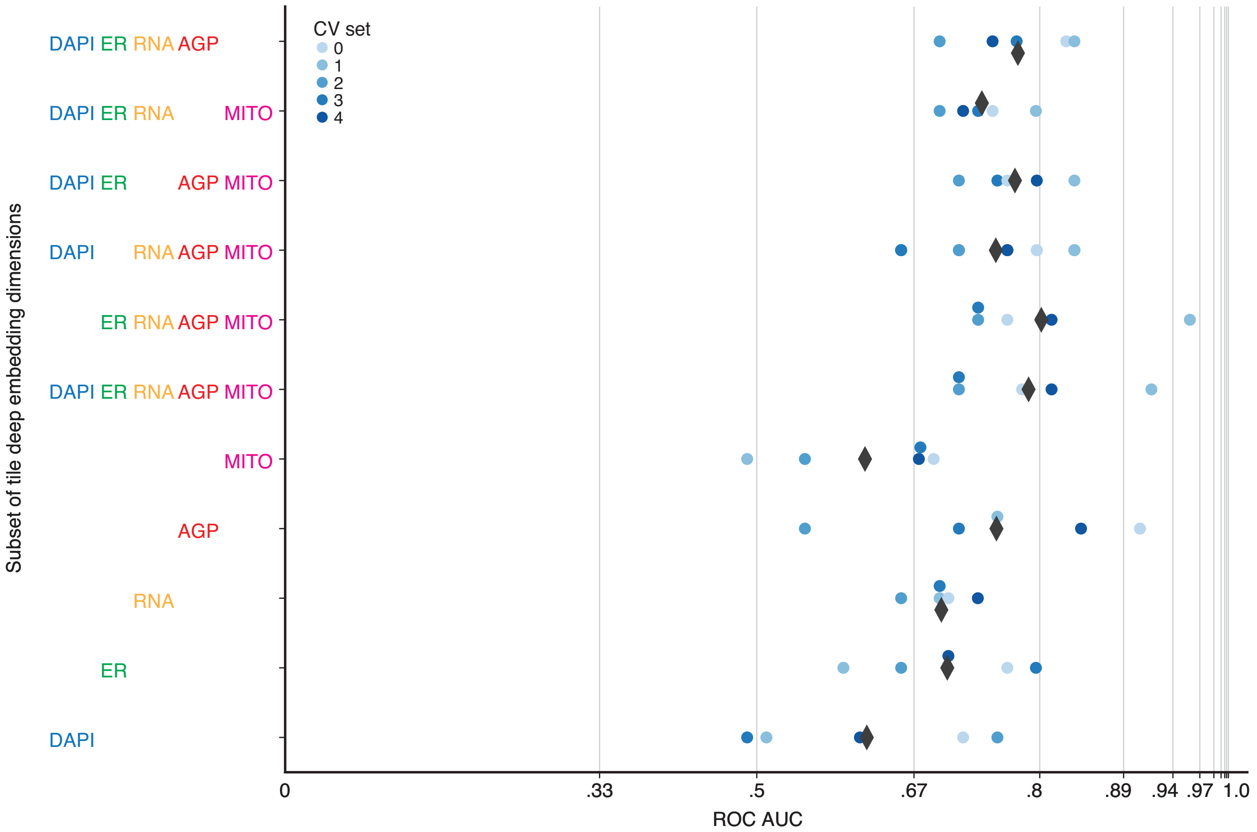


**Supplementary Fig. 5 | Impact of individual Cell Painting channels on PD classification.** The same logistic regression model with tile deep embeddings from **Fig. 5b** evaluated with a subset of the deep embedding dimensions corresponding to a subset of the 5 channels. Black diamonds denote the mean across all cross-validation (CV) sets. Grid line spacing denotes a doubling of the odds of correctly ranking a random healthy control and PD cell line. Dashed line denotes chance performance.

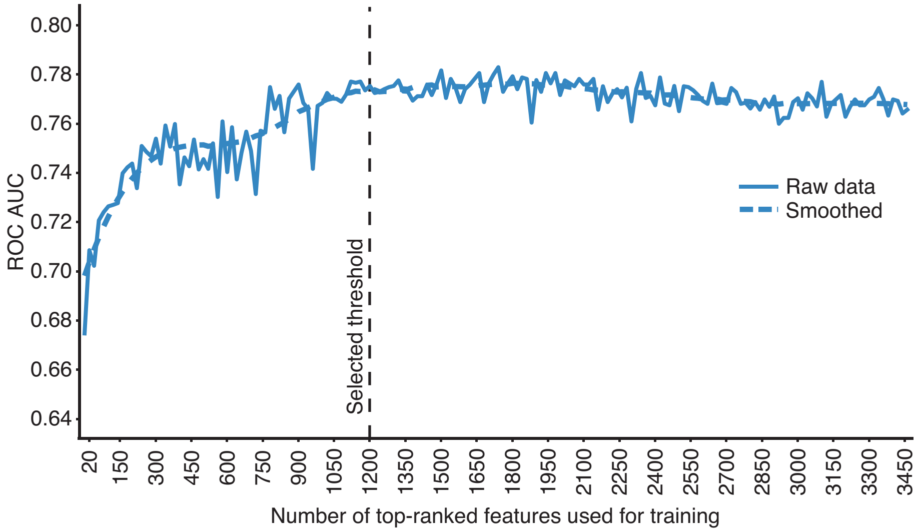


**Supplementary Fig. 6 | Estimating threshold for number of top-ranked CellProfiler features required for PD classification.** Performance of the random forest classifier as a function of number of top-ranked features used for training, evaluated in increments of 20 features. The dashed line represents the threshold selected for subsequent analyses.
